# Self-healing of hyaluronic acid to improve *in vivo* retention and function

**DOI:** 10.1101/2021.09.17.460792

**Authors:** Anna Gilpin, Yuze Zeng, Jiaul Hoque, Ji Hyun Ryu, Yong Yang, Stefan Zauscher, William Eward, Shyni Varghese

## Abstract

Convergent advances in the field of soft matter, macromolecular chemistry, and engineering have led to the development of biomaterials that possess autonomous, adaptive, and self-healing characteristics similar to living systems. These rationally designed biomaterials could surpass the capabilities of their parent material. Herein, we describe the modification of hyaluronic acid (HA) molecules to exhibit self-healing properties and studied its physical and biological function both *in vitro* and *in vivo*. Our *in vitro* findings showed that self-healing HA designed to undergo autonomous repair improved lubrication, enhanced free radical scavenging, and resisted enzymatic degradation compared to unmodified HA. Longitudinal imaging following intra-articular injection of self-healing HA showed improved *in vivo* retention despite the low molecular weight. Concomitant with these functions, intra-articular injection of self-healing HA mitigated anterior cruciate ligament injury-mediated cartilage degeneration in rodents. This proof-of-concept study shows how incorporation of functional properties like self-healing can be used to surpass the existing capabilities of biolubricants.

## Introduction

Incorporating distinct molecular and chemical features into biomaterials can introduce new functionalities to combat intrinsic limitations of the parent materials towards their desired application. For example, biomaterials that can undergo *in situ* self-repair could not only improve their *in vivo* longevity and function but could also promote their therapeutic outcome. Herein, we examine whether hyaluronic acid molecules designed to undergo self-healing could improve their physical and biological functions with a focus on biolubrication. Biolubrication is critical for the efficient function of diarthrodial joints, eyes, lungs and other visceral organs. The lubricating molecules present at the articular cartilage interfaces of the diarthrodial joints maintain tissue health and tribological functions.^[1]^ Enabling inter-surface lubrication is also important for the seamless function of various medical devices. Due to its prevalence in biolubrication and biological functions^[2]^ and its alteration in diseases like osteoarthritis,^[3]^ HA has been widely used to treat lubrication deficiencies.^[4]^ Besides promoting lubrication, HA also exhibits various biological functions, such as reducing inflammation and free radical damage, as well as alleviating pain.^[5]^ However, a persistent challenge to HA lubricants is their short *in vivo* residence time.^[6]^ Hence, strategies that ensure long-term retention and function of exogenous HA have been developed to improve its clinical outcome.^[7]^ One of the most widely used approaches to enhance the *in vivo* longevity of HA includes the introduction of chemical crosslinks; however, this strategy severely limits HA’s lubricating function and its handling.^[5a, 8]^ Albeit at low incidence, chemical crosslinking can also result in pseudoseptic reactions.^[9]^

We introduced self-healing and dynamic crosslinking into HA molecules and assessed their potential to improve the physical and biological functions of parent HA without compromising its injectability. Towards this, we modified HA with ureidopyrimidinone (UPy) groups to enable self-healing of HA polymers under physiological conditions, *via* reversible secondary interactions.^[10]^ Biomaterials modified with UPy molecules have been exploited for various biomedical applications.^[11]^ UPy molecules rapidly dimerize through quadruple hydrogen bonding resulting in dynamic supramolecular structures under physiological conditions.^[10, 12]^ HA molecules endowed with UPy moieties can form a dynamic network through hydrogen bonding while exhibiting shear-thinning behavior (*via* reorganization of the polymer chains in response to shear forces), thus enabling easy injection and efficient lubrication. At rest, the rapid UPy dimerization re-establishes the stable supramolecular network, and these “self-generating” networks will resist rapid clearance from the synovial space (**Fig. 1a**). Thus, the self-healing HA molecules will offer the benefits of both high molecular weight HA (shear thinning, mechanical adaptability, and enhanced lubrication), as well as chemically crosslinked HA (improved *in vivo* retention and reduced enzymatic degradation). As a proof-of-concept, we modified 200 kDa HA molecules with UPy moieties and assessed whether self-healing capability could improve the *in vivo* retention and chondroprotective function of HA in articular cartilage lubrication. We chose to use low molecular weight HA because it lacks viscoelasticity and other beneficial effects towards *in vivo* retention, thus eliminating any confounding contributions.

**Figure 1.**
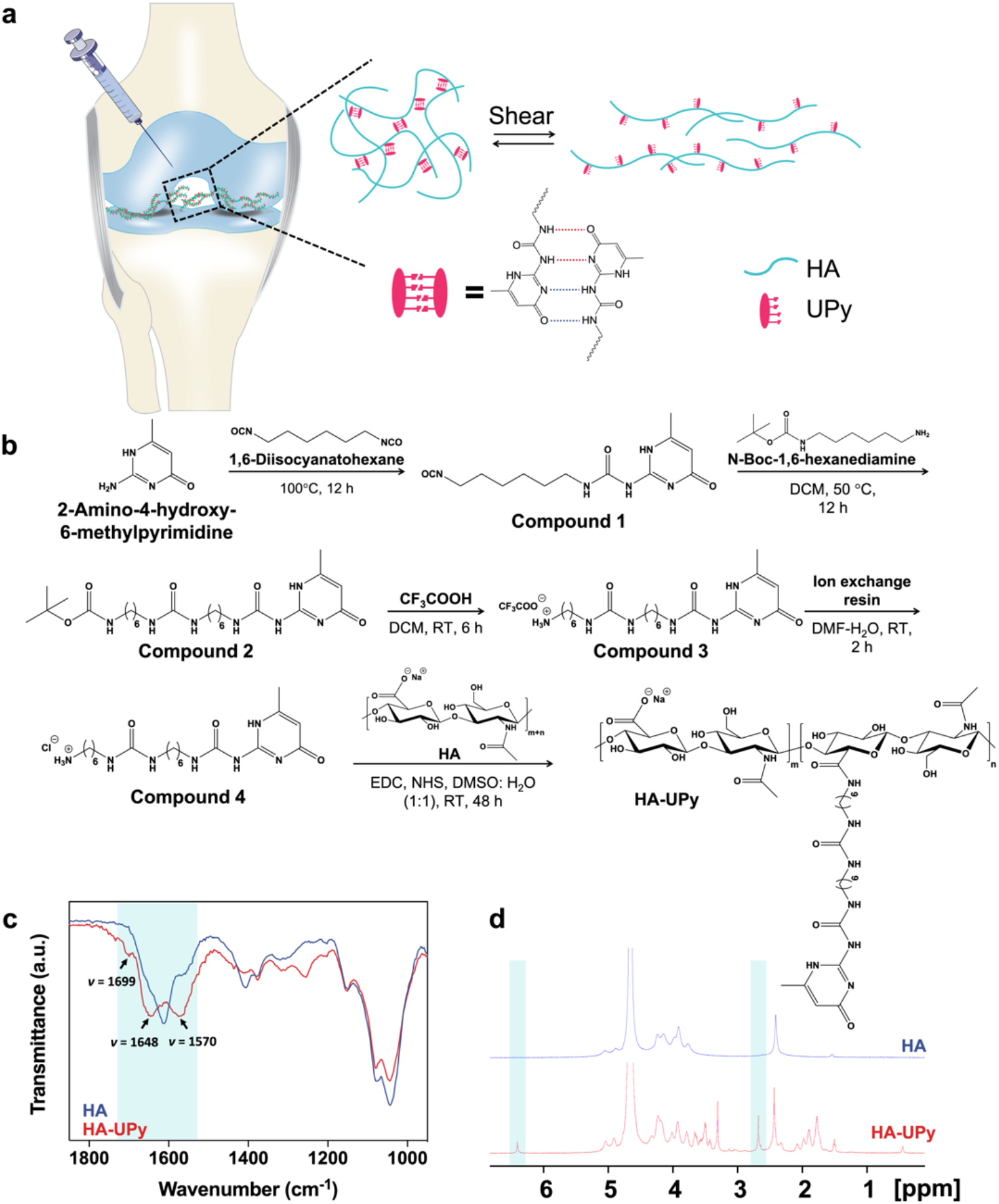
Development of self-healing HA lubricant (i.e, HA-UPy). ***a***, Schematics of intra-articular injection of self-healing HA lubricant. When under shear during injection or joint movements, HA-UPy becomes fluidic and spreads in the joint, lubricating the articular surface. After the removal of shear force, HA-UPy can self-generate into a network *via* UPy-mediated hydrogen bonding, contributing to its retention in the joint. ***b***, Synthesis route. DCM: dichloromethane; DMF: dimethylformamide; DMSO: dimethyl sulfoxide; EDC: 1-ethyl-3-(3-dimethylaminopropyl) carbodiimide hydrochloride; NHS: N-hydroxysuccinimide; RT: room temperature. ***c***, FTIR spectra of HA and HA-UPy. ***d***, ^1^H NMR spectra. Pyrimidinone protons of UPy: δ 6.41 ppm (=CH−) and δ 2.69 ppm (−CH_3_).

## Results and discussion

### Synthesis and characterization of HA-UPy molecules

HA-UPy molecules were prepared by functionalizing HA with UPy-bearing linkers as shown in the reaction scheme (**Fig. 1b)**. Details of the UPy linker synthesis are provided in Methods and Supplementary Information (**Supplementary Figs. 1-5 and Supplementary Note**). The linker was designed to bear the UPy moiety on one end^[12a]^ with the other end containing a primary amine group protected by a cleavable tert-butyloxycarbonyl (Boc) group. Upon removal of the Boc group, the primary amine on the linker reacts with the carboxylic acid of the HA’s D-glucuronic acid, resulting in UPy conjugation to HA. The Fourier-transform infrared (FTIR) spectroscopy of the product confirmed UPy conjugation with new peaks appearing at 1699, 1648, and 1570 cm^-1^, corresponding to pyrimidinone C=O stretching, urea C=O stretching, and pyrimidinone C=N stretching, respectively (**Fig. 1c**). The UPy conjugation was further confirmed by proton nuclear magnetic resonance (^1^HNMR) spectroscopy (**Fig. 1d**). We estimated a grafting density of ∼24 ± 3% per dimeric repeating unit of HA from the ^1^HNMR spectra by comparing the integrated peak area of the pyrimidinone protons in a UPy unit (−NH−C(CH_3_)−C<u>H</u>−CO−, 1H, δ 6.41; −NH−C(<u>CH_3_</u>)−CH−CO−, 3H, δ 2.69) to that of the acetyl protons in HA’s N-acetyl-D-glucosamine unit (−NH−CO−<u>CH_3_</u>, 3H, δ 2.45) (**Fig. 1d and Supplementary Fig. 6**). For live *in vivo* imaging (IVIS), HA and HA-UPy molecules were conjugated with the near infrared (NIR) fluorescent dye cyanine 7 (Cy7) (**Supplementary Fig. 7)**.

### HA-UPy molecules form supramolecular networks and exhibit self-healing

The UPy-mediated non-covalent interactions among the polymer chains can facilitate self-assembly of HA molecules into dynamic networks (*i*.*e*., soft gels), which we characterized by rheological measurements. To study the effect of polymer concentration, we prepared solutions of HA-UPy and HA at three concentrations (2, 5, and 10 wt%) in phosphate-buffered saline (PBS). The frequency sweep (0.1–10 Hz) measurements of HA-UPy and unmodified HA showed that HA-UPy samples exhibit higher G’ (storage modulus) and G” (loss modulus) at all frequencies while the HA samples behaved more like a viscous liquid^[13]^ (**Fig. 2a and Supplementary Figs. 8a, b**). Furthermore, the storage modulus, determined at 1 Hz, of the HA-UPy samples increased with increasing concentration (**Fig. 2b**). The oscillation frequency of 1 Hz was chosen because it is within the range of typical walking frequencies.^[14]^ The concentration-dependent network formation was also visualized by inverting the tubes containing the polymer solutions. The samples containing HA-UPy showed solid-like behavior at higher concentrations and did not flow like a liquid. In contrast, the samples containing unmodified HA behaved like a viscous liquid at all concentrations, with the 10 wt% solution taking a longer time to flow (**Supplementary Fig. 8c**). These observations for the HA-UPy samples are consistent with network formation, which arises from quadruple hydrogen bonding between the UPy moieties. Further experiments were carried out using 10 wt% HA and HA-UPy.

**Figure 2.**
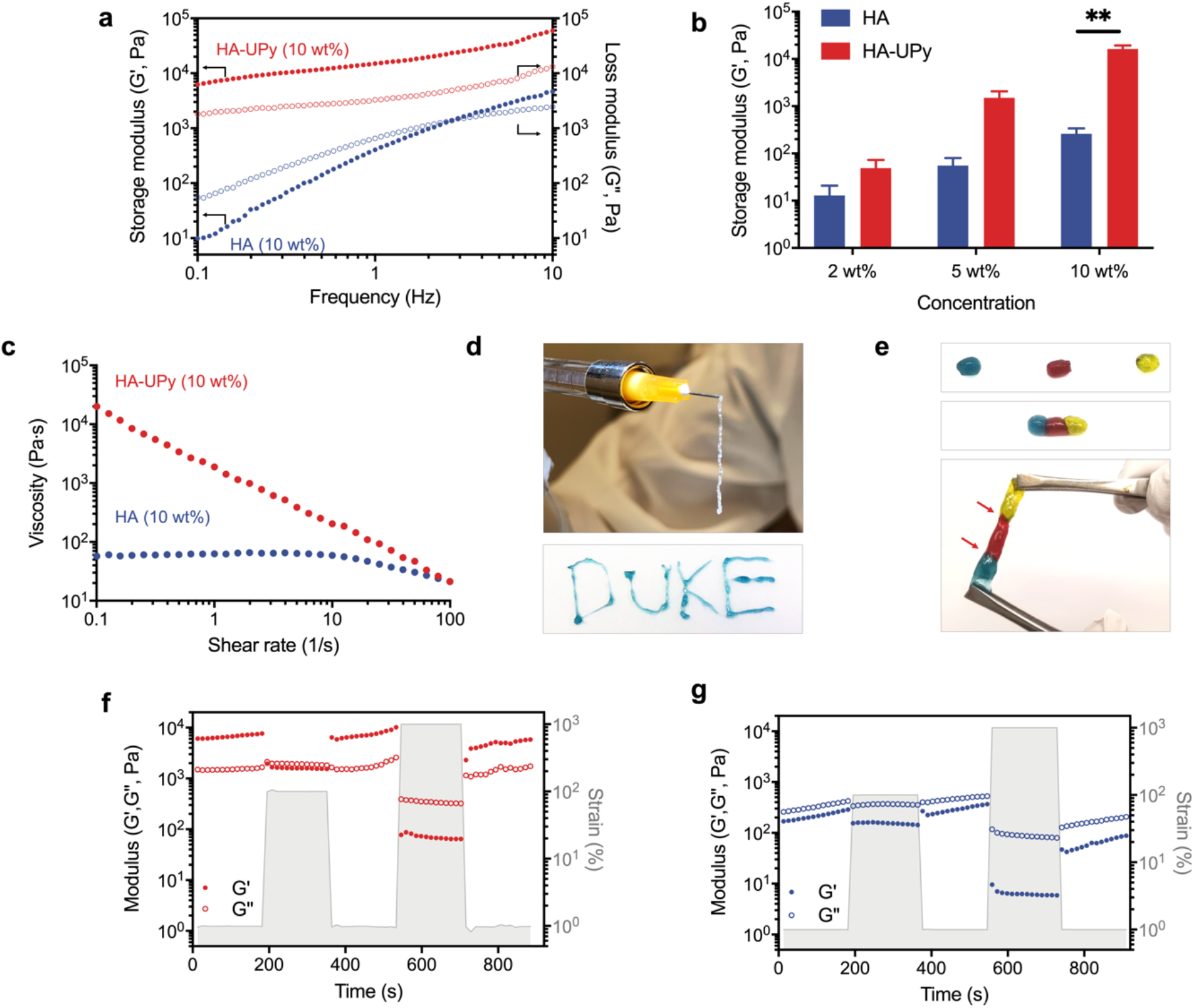
Characterization of self-healing HA. ***a***, Storage (G′) and loss (G′′) moduli of HA-UPy and HA in a frequency sweep measurement. ***b***, Storage modulus (G′) of HA-UPy and HA as a function of concentration. ***c***, Viscosity changes of HA-UPy and HA as a function of shear rate. ***d***, HA-UPy (10 wt%) was easily extruded through a 26G needle into “DUKE” letters. ***e***, Separate pieces of HA-UPy (10 wt%) hydrogels healed together with interfaces indicated with red arrows. ***f***,***g***, Step-strain measurements for 10 wt% HA-UPy (**f**) and 10 wt% HA (**g**).

Because the UPy-mediated network formation is dynamic, the HA-UPy samples should show shear-thinning and self-healing functions. As expected, the viscosity of the HA-UPy samples decreased with increasing shear rate, showing a characteristic shear-thinning behavior which results from the destruction of the physical crosslinks by the applied shear stresses (**Fig. 2c**). In contrast, the shear rate-dependent viscosity of the corresponding HA solution is consistent with that of a viscous liquid. The HA-UPy samples (10 wt%) were easily injected, with minimal resistance, through a 26G hypodermic needle (**Fig. 2d**). The extruded HA-UPy formed a stable network at rest, which enabled the “printing” of different shapes (**Fig. 2d**).

We examined the self-healing of HA-UPy, following an approach reported previously,^[15]^ by bringing multiple pieces of HA-UPy hydrogels into close contact, which showed instantaneous healing (**Fig. 2e**). Furthermore, we used step-strain measurements to confirm the UPy-mediated self-healing of HA-UPy and dynamic networks, wherein 10 wt% HA-UPy and HA samples were subjected to alternating step strains of 1 to 100% and 1 to 1000% (**Figs. 2f, g)**. The storage modulus (G’) values of the HA-UPy samples dropped to that of the loss modulus (G”) at a strain γ =100%, indicating network disruption (**Fig. 2f**). When the strain was removed, the HA-UPy molecules re-organized and formed a new network structure instantaneously with a 100% recovery of G’. Increasing the strain rate to 1000% induced more network destruction as indicated by the drop of the storage modulus to ∼100 Pa, with a corresponding inversion of G′ and G″, suggesting liquid-like flow behavior. Despite the large strain (γ = 1000%), prompt recovery of the network structure was observed upon the removal of the strain. Unmodified HA samples, on the other hand, had a higher loss modulus than storage modulus at both low and high strains (**Fig. 2g**). This behavior is indicative of a viscous liquid. Additionally, we performed experiments where we alternatingly applied a low (1%) and high (500%) strain over multiple cycles to determine whether the HA-UPy sample would recover its storage modulus after repeated network disruptions. These studies showed complete network formation without hysteresis as indicated by the G′, which maintained its original value at 1% strain following repeated network disruption at γ =500% (**Supplementary Fig. 9**).

### Self-healing HA exhibits enhanced lubrication

Effectiveness of the biolubricant to reduce friction between the articular surfaces is key to its application in improving joint function. We thus investigated whether UPy-mediated changes in the viscoelastic properties of HA-UPy would translate to its lubrication function. To this end, we determined the coefficient of friction (µ) between healthy porcine cartilage explants in the presence of HA-UPy and compared the measured values with those obtained when using corresponding HA solutions and saline (negative control) by using a rotational rheometer. We determined the coefficient of friction (COF), µ, at the cartilage-to-cartilage interface, using the equation, 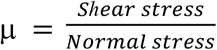 **Figure 3a** shows that the COF between the contacting articular surfaces decreased significantly in the presence of HA-UPy. Specifically, the HA-UPy molecules reduced friction by ∼70% and 55% compared to saline and HA molecules, respectively.

**Figure 3.**
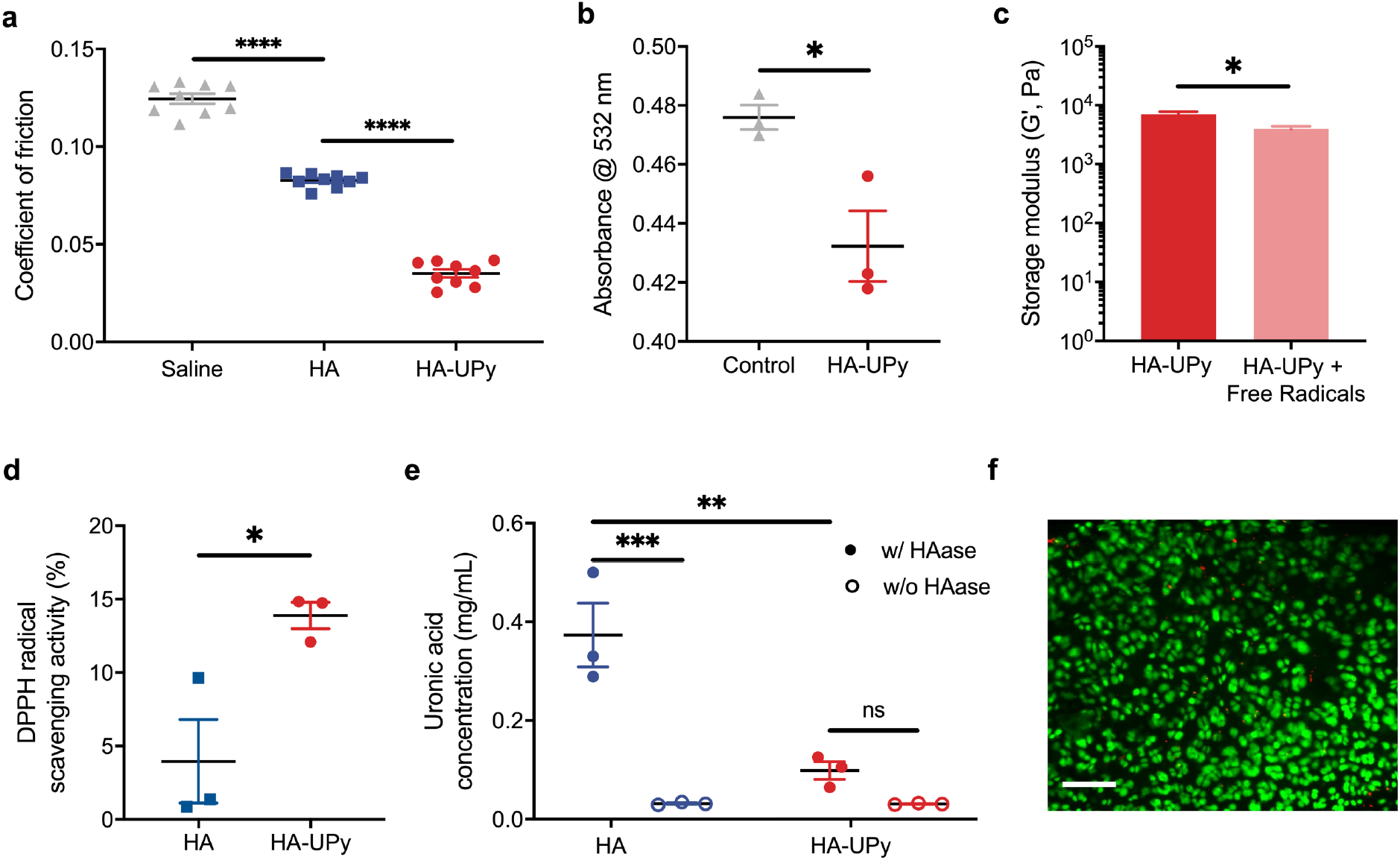
Lubrication, antiradical capacity, *in vitro* stability, and cytocompatibilty of self-healing HA. ***a***, Coefficients of friction at the cartilage-cartilage interface in the presence of HA-UPy, HA, and saline. ***b***, Free radical scavenging effect of HA-UPy exposed to Fenton reagent compared to the phosphate buffer control. ***c***, Storage modulus (G′) of HA-UPy measured at a frequency of 1 Hz following exposure to hydroxyl radicals compared to non-exposed HA-UPy. ***d***, DPPH radical scavenging of HA-UPy compared to HA. ***e***, Degradation products of HA-UPy and HA with and without the presence of hyaluronidase (HAase). ***f***, Chondrocyte viability in a rat cartilage explant after 7 d incubation with HA-UPy. Green: live cells; Red: dead cells. Scale bar: 100 μm. All data are presented as means (± s.e.m.). One-way ANOVA ***(a)***, unpaired two-tailed t-test ***(b, c, d)*** or two-way ANOVA ***(e)*** with Tukey’s multiple-comparisons test was used for statistical analysis. Significance is determined as \**P* < 0.05, \*\**P* < 0.01, \*\*\**P* < 0.001, \*\*\*\**P* < 0.0001 and n.s. (not significant).

### Self-healing HA promotes free radical scavenging

HA has a number of biological functions, including serving as an antioxidant to reduce free radical damage to cells. We investigated the free radical scavenging effect of HA-UPy using deoxyribose/Fenton reagent and 1,1-diphenyl-2-picrylhydrazyl (DPPH) assays.^[16]^ In the deoxyribose/Fenton reagent assay, hydroxyl radicals are produced by the reaction of Fe^2+^−EDTA with hydrogen peroxide. The hydroxyl radicals subsequently interact with deoxyribose and form a pink color chromogen with thiobarbituric acid upon heating. Following incubation with the Fenton reagent, the absorbance of the solution was measured. As seen in **Figure 3b**, the solution incubated with HA-UPy had a lower chromogen absorbance, corresponding to fewer hydroxyl radicals, suggesting free radical scavenging by HA-UPy molecules. Since free radicals cleave the glycosidic bonds in HA which could lead to the breakdown of polymer chains, we examined the storage moduli of the HA-UPy exposed to the Fenton reagent and compared them to the storage moduli of untreated HA-UPy samples. **Figure 3c** shows a slight reduction in the G′ value, indicating some disruption of the network in the presence of free radicals (**Supplementary Fig. 10**). Although we stopped the reaction after 1 h, ensuring the complete removal of free radicals from the solution is challenging. It is thus likely that free radicals continued to react with HA-UPy molecules. While the Fenton assay enables us to examine the free radical scavenging effect of the HA-UPy in a physiologically relevant environment, a potential radical scavenging property of unmodified HA molecules cannot be determined because of the aqueous reaction environment.

Hence, we also used a DPPH assay to examine the UPy-mediated changes in free radical scavenging, where the samples were exposed to a DPPH solution in ethanol. The solution containing HA-UPy molecules showed a significantly reduced DPPH free radical concentration as compared to that containing HA, which had a minimal scavenging effect (**Fig. 3d**). The free radical scavenging ability of HA-UPy could be due to the network formation and/or the presence of UPy moieties. Prior studies have showed that the protective effect of HA against the free radical damage to the cells depends on HA molecular weight, with high molecular weight HA providing better protection.^[17]^ Furthermore, UPy moieties contain pyrimidine rings, which are known to scavenge free radicals.^[18]^ The minimal reduction in G′ of HA-UPy following free radical exposure could be attributed to the UPy-mediated self-healing/self-generation of networks or by the UPy scavenging the free radical itself.

### Self-healing HA molecules resist enzymatic degradation

HA within the synovial fluid is subjected to enzymatic and free radical degradation as well as lymphatic drainage, which are some of the key players contributing to its rapid clearance in the joint. The short residence time (t_1/2_ ∼ 24 h) of HA within the synovial joint^[19]^ has been thought to be one of the factors contributing to its limited clinical effectiveness following intraarticular injection, and chemically crosslinked HA derivatives have thus been generated to delay or slow the breakdown.^[5a, 20]^ We posit that the formation of supramolecular HA networks by UPy interactions may also slow the degradation of HA molecules. To test this, we incubated HA-UPy with hyaluronidase and quantified the resultant HA fragments by using a modified uronic acid assay.^[21]^ As demonstrated by the results, the HA-UPy experienced minimal degradation compared to the corresponding HA in the presence of hyaluronidase. Moreover, no statistical significance is observed between HA-UPy incubated with hyaluronidase and controls (*i*.*e*., HA and HA-UPy in the absence of hyaluronidase) (**Fig. 3e**). The diminished degradation of HA-UPy is likely due to the UPy-mediated network formation, which could shield the enzyme-specific binding sites from hyaluronidase.^[21]^

Given the direct contact between HA-UPy and the cartilage surface, we also evaluated the cytocompatibility of HA-UPy by exposing rat cartilage explants to HA-UPy for a duration of 7 days. The live/dead analyses showed that nearly 100% of the chondrocytes were alive with no detrimental effect (**Fig. 3f**).

### Self-healing improved the *in vivo* retention of HA

The *in vivo* retention of HA-UPy following intraarticular injection was studied as a function of time by live imaging of rat knees and compared against corresponding HA. Cy7-conjugated HA-UPy and HA molecules were injected into rat knees and monitored using IVIS for 28 days. We performed calibration studies to ensure that both cohorts received similar levels of Cy7 molecules. The animals were imaged immediately and 24 h post-injection for the initial reading, which showed clear positive signals from the joints administered with the HA-UPy and HA molecules. Longitudinal imaging indicated that a majority (>60%) of the HA was cleared from the joint by ∼3 d (**Figs. 4a, b**). In contrast, strong positive signals were present in the HA-UPy group even at 28 d, the maximum experimental time point, with a ∼40% reduction in fluorescence intensity compared to the initial reading (*i*.*e*., immediately after administration) (**Fig. 4b**). The values are presented as a percentage of initial fluorescence intensity to account for variability among the animals.

**Figure 4.**
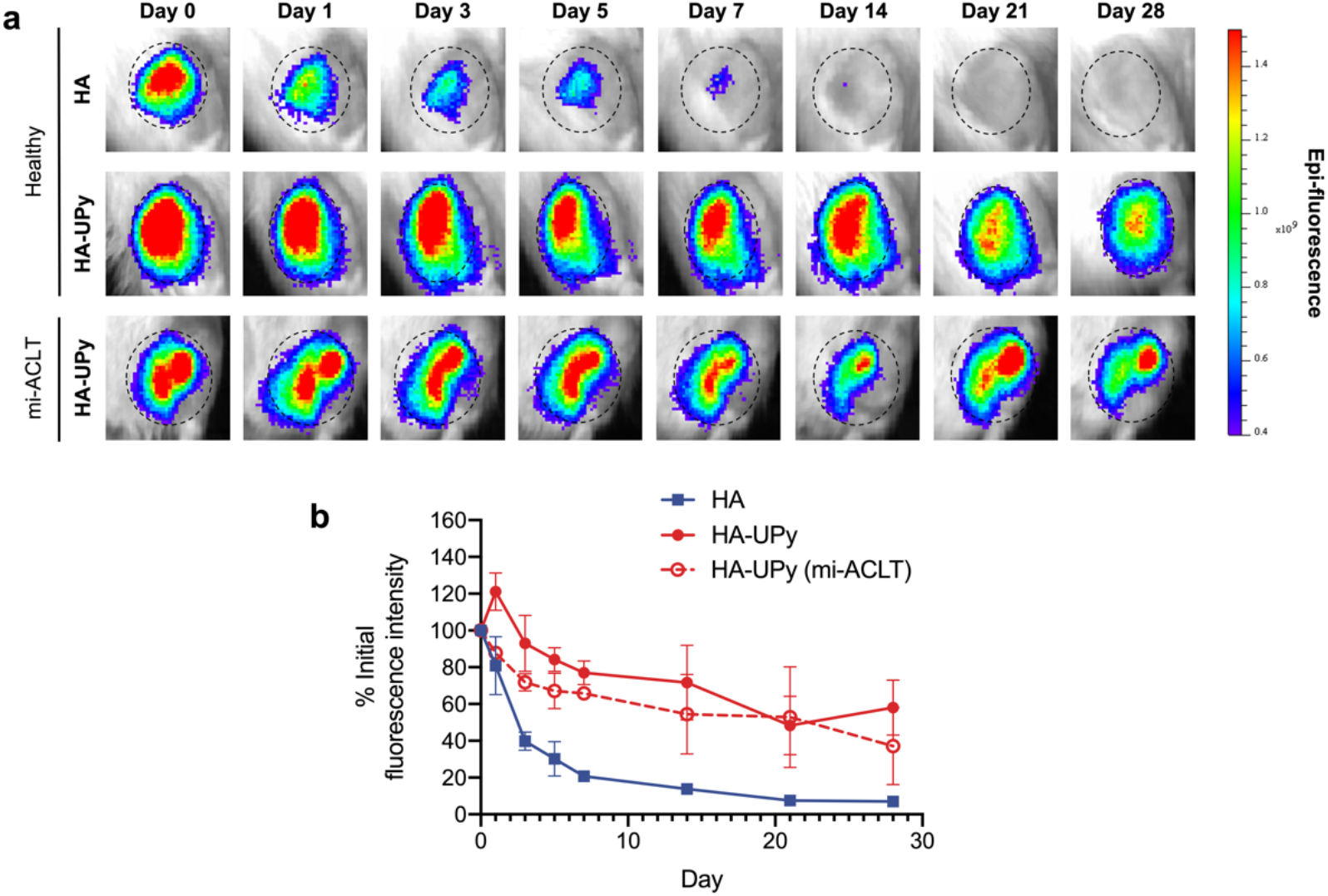
*In vivo* retention of self-healing HA. ***a***, Representative IVIS images of rat knee joints following intra-articular injection of Cy7-tagged HA or HA-UPy as a function of time. Dashed circles demarcate the joint region of interest (ROI) that was used for fluorescence intensity quantification. Color map reflects the epi-fluorescence intensity with red being the strongest. ***b***, Quantification of fluorescence intensity of ROI as a percentage of initial intensity. HA: n=2, HA-UPy: n=3, HA-UPy (mi-ACLT): n=2. Data are presented as means (± s.e.m.).

The effect of injury on lubricant clearance was examined by comparing the retention of HA-UPy in rat joints which underwent minimally invasive anterior cruciate ligament transection (mi-ACLT), where the HA-UPy molecules were administered two days post-mi-ACLT. Similar to the healthy group, a significant amount of the administered HA-UPy was retained within the mi-ACLT joints, albeit less than that in the uninjured joints. Given that a molecular weight of 200 kDa is well below the permeability barrier of the synovial membrane,^[22]^ we attribute the increased residence time of HA-UPy within the joint synovial space to self-healing of HA molecules.

### Self-healing HA provides improved chondroprotection

The enhanced lubrication along with its improved retention in the joint suggests that the self-healing HA-UPy could offer chondroprotection following joint injury. To assess the *in vivo* chondroprotective function, we have used mouse and rat ACL transection models. The surgical ACL transection model is widely used to represent articular cartilage degeneration consistent with ACL injuries, which cause joint instability, chronic inflammation, and degeneration.^[23]^ The ACL-transected mice received weekly intraarticular injections of HA-UPy, HA, or saline for four weeks beginning one week post-surgery as shown in the experimental timeline (**Supplementary Fig. 11a**). Weekly injections were chosen based on prior reports^[24]^ and *in vivo* imaging which showed complete clearance of HA by day 7. Safranin O staining of the knee joints at week 5 showed significant damage to the articular surfaces of the cohorts that received saline (**Supplementary Fig. 11b**). Similar to the saline group, the animals that received HA injections showed significant cartilage degeneration. In contrast, the cohort that received HA-UPy maintained better cartilage integrity with significant positive staining for glycosaminoglycans. We used a semi-quantitative score of cartilage degeneration (OARSI) to assess the matrix loss, surface fibrillation, and erosion in the joint, which corroborated the histological findings (**Supplementary Fig. 11c**). The lower scoring value for the animals that received HA-UPy suggests improved chondroprotective function of the parent HA following modification (i.e., HA-UPy).

Because mouse joints only permit intraarticular injections of small volumes (∼5 μL), we employed a rat knee injury model to determine the chondroprotection of self-healing HA-UPy. By using a minimally invasive, percutaneous procedure, we have developed an ACL injury (mi-ACLT) in the rat knee without surgically opening the joint. Specifically, the ACL was transected with an 18G needle which was inserted into the knee joint lateral to the patellar ligament while the knee was flexed at a ∼120° angle (**Supplementary Fig. 12a**). Successful ACL rupture was confirmed by using the anterior drawer test, which exhibited abnormal subluxation of the tibia. The dissected knee joints post-mortem showed that the ACL had been successfully transected (**Supplementary Fig. 12b**). The mi-ACLT-mediated cartilage degeneration was assessed at week 9 following weekly saline injections over eight weeks. Safranin O/Fast Green staining of sagittal sections of the articular joints showed severe fibrillation and erosion of both the tibial and femoral cartilage (**Supplementary Fig. 12c**). The formation of osteophytes was also visible on the posterior region of the tibia. In contrast, the cartilage surfaces of the uninjured contralateral limbs were smooth with no significant degeneration, and no osteophytes were present. The degree of degeneration was quantified using the rat OARSI score,^[25]^ which showed that the mi-ACLT group had a significantly higher score than that of the unoperated control, consistent with greater cartilage degeneration (**Supplementary Fig. 12d**).

The extent of cartilage degeneration was further examined by immunohistochemical (IHC) staining for catabolic markers— matrix metalloproteinase-13 (MMP-13) and a disintegrin and metalloproteinase with thrombospondin motifs 5 (ADAMTS-5), which are shown to be highly active during cartilage degeneration.^[26]^ Cartilage in the mi-ACLT group showed a higher expression of both ADAMTS-5 and MMP-13 than the contralateral group, indicating greater degeneration (**Supplementary Fig. 12e**). Furthermore, clustering of the chondrocytes, a hallmark of osteoarthritic cartilage,^[27]^ was clearly present in the mi-ACLT group but not in the healthy contralateral group. In addition to the semi-quantitative scoring (OARSI score), total cartilage degeneration (matrix, proteoglycan, or chondrocyte loss), significant cartilage degeneration (degeneration >50% of cartilage thickness), surface matrix loss (matrix fibrillation), and the depth ratio of cartilage lesions (ratio of depth of cartilage degeneration to total cartilage thickness, measured at three zones) were quantified as described elsewhere.^[25]^ In addition to the tibia (which is commonly the focus for rat OARSI scoring), the femur was also analyzed, as recent studies have shown that the medial femoral condyle exhibits severe degeneration with ACL injury, both in animals^[28]^ and humans.^[29]^ The mi-ACLT was shown to significantly increase total degeneration (**Supplementary Fig. 12f**), significant cartilage degeneration (**Supplementary Fig. 12g**), surface matrix loss (**Supplementary Fig. 12h**), and depth ratio of cartilage lesions (**Supplementary Fig. 12i**) as compared to the healthy contralateral group. Together the data demonstrate significant cartilage degeneration following mi-ACLT.

We next examined the chondroprotective function of self-healing HA in the rat mi-ACLT model. Because the HA- and saline-treated animals exhibited similar cartilage degeneration, we have compared the HA-UPy-treated rat joints to those treated with corresponding HA. As described in **Fig. 5a**, the animals received weekly injections starting one day post-mi-ACLT for a total of eight weeks. At week 9, animals were euthanized, and their joints were examined histologically. The cohorts treated with HA showed significant cartilage degeneration compared to those treated with HA-UPy. Specifically, cartilage erosion, proteoglycan loss, and osteophytes in the tibia were clearly observed in the HA group and were similar to features observed in the saline group (**Fig. 5b**). On the contrary, joints treated with HA-UPy showed higher Safranin O staining intensity with minimal cartilage thinning but displayed some degree of cartilage fibrillation and osteophyte formation. In comparing the OARSI scores, HA-UPy, while higher than the contralateral group, had a significantly lower score than the corresponding HA group (**Fig. 5c**).

**Figure 5.**
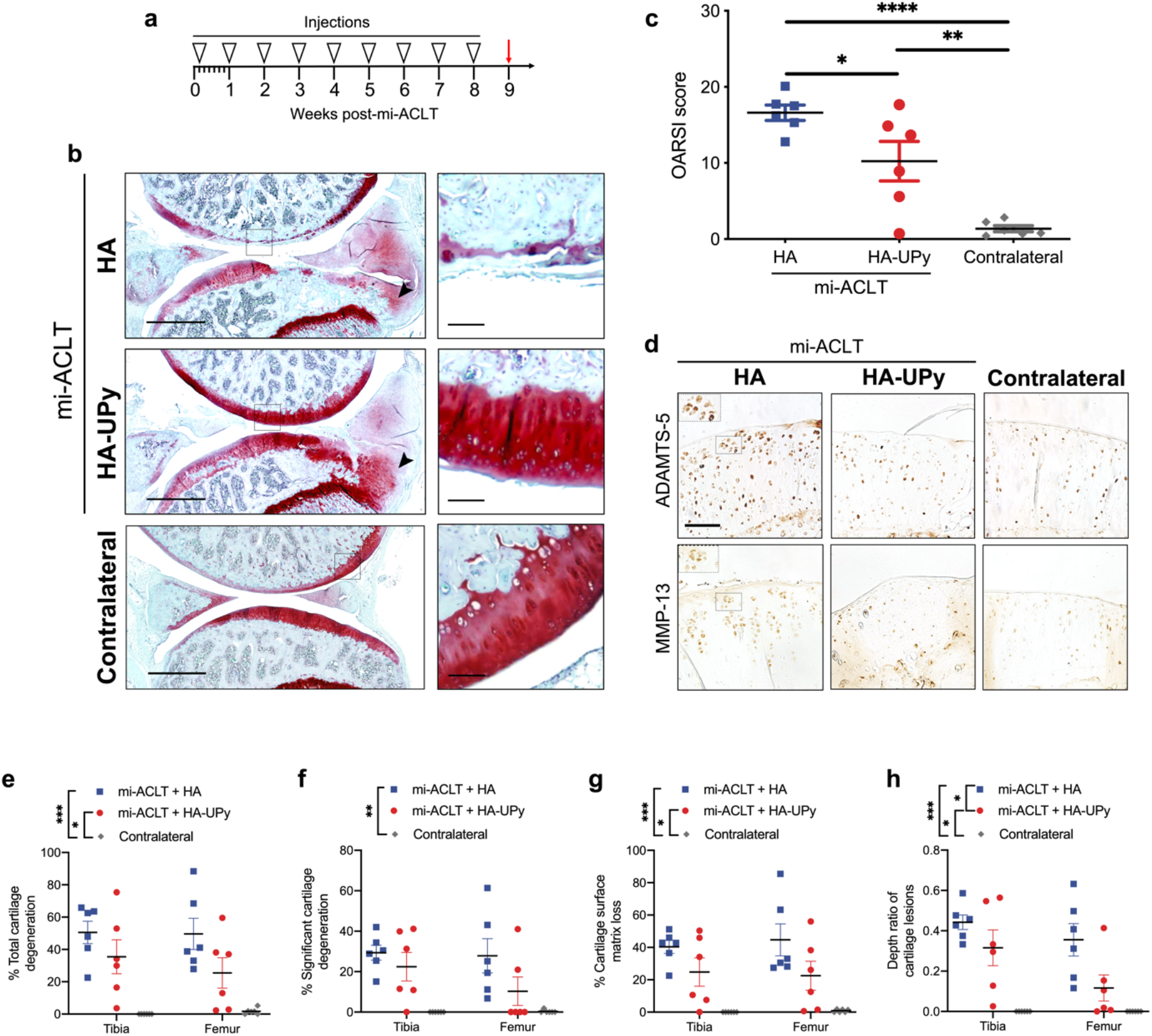
Chondroprotection of self-healing HA in a minimally invasive rat ACLT model. ***a***, Experimental timeline showing schedule of injections. ***b***, Safranin O-stained mi-ACLT joint treated with HA showed severe cartilage degeneration with an osteophyte in the tibia (arrow). HA-UPy-injected joints showed strong proteoglycan staining while exhibiting some cartilage fibrillation and osteophyte formation (arrow). Contralateral joints without injury were used as a positive control. Scale bar: 1 mm. ***c***, OARSI scoring indicates that the HA-UPy group had significantly less degeneration than the unmodified HA group. Data are presented as means (± s.e.m.) and statistical significance was analyzed using one-way ANOVA with Tukey’s multiple comparisons test. ***d***, ADAMTS-5 and MMP-13 IHC staining of tibial cartilage. Greater positive staining and chondrocyte clustering (inset) is observed in the HA group compared to the HA-UPy group. Scale bar: 100 μm. ***e-h***, Quantitative measures of ***(e)*** total cartilage degeneration, ***(f)*** significant cartilage degeneration, ***(g)*** cartilage surface matrix loss, and ***(h)*** thickness of cartilage lesions for both the tibia and femur. Data are presented as means (± s.e.m.). A two-way repeated measures ANOVA was used to analyze statistical significance with Tukey’s multiple comparisons used to analyze the differences between treatments. The significance in the legend shows the Tukey’s multiple comparisons between treatments. Significance is determined as \**P* < 0.05, \*\**P* < 0.01, and \*\*\**P* < 0.001.

Furthermore, MMP-13 and ADAMTS-5 IHC staining showed higher expression of these catabolic enzymes in the HA group, as seen by greater staining (both in intensity and the number of stained cells), compared to the HA-UPy and contralateral groups (**Fig. 5d**). Furthermore, the joints treated with HA showed evidence of chondrocyte clustering, similar to those treated with saline. A majority of the HA-UPy-treated joints showed minimal cartilage degeneration, and no chondrocyte clustering was observed in these animals similar to the unoperated contralateral groups. Moreover, the organization and distribution of chondrocytes within the cartilage of cohorts treated with the HA-UPy molecules was found to be similar to that of the uninjured contralateral groups. We also quantitively assessed the total cartilage degeneration, significant cartilage degeneration, surface matrix loss, and the depth ratio of cartilage lesions. These parameters were lower in the HA-UPy group than the HA group for both the femur and tibia (**Figs. 5e-h**). Despite the high variability among the treated animals, the total tibial degeneration was ∼30% less in animals treated with HA-UPy compared to those treated with HA (as a percentage of total cartilage width: 35±10% for HA-UPy *vs*. 51±7% for HA). Similarly, joints treated with HA-UPy displayed half as much total cartilage degeneration in the femur (as a percentage of total cartilage width: 25±9% for HA-UPy *vs*. 50±10% for HA) compared to those treated with HA. The HA-UPy group also showed a reduced amount of significant cartilage degeneration, which consists of the width of cartilage in which 50% or more of the cartilage thickness is degenerated. The cartilage lesions in the HA-UPy group also spanned significantly less of the cartilage thickness as compared to those in the HA group were also significantly less deep than the HA group, indicating that self-healing HA was more chondroprotective than unmodified HA. The high variability observed in the HA-UPy-treated group could be attributed to the presence of minimally modified or unmodified HA molecules. It is also likely that the high variability is due to differences in the initial cartilage damage that may result from the needle during the mi-ACLT procedure. Because the injury is performed on a closed knee, there is risk of unintentionally damaging the cartilage or other joint tissues, increasing the severity of the injury. While the potential for this variability is high, we have randomized the animals to each treatment to ensure that differences due to the mi-ACLT procedure are spread amongst groups.

## Conclusion

Taken together, our results show that endowing HA with self-healing functionality can be used to improve the intraarticular longevity of HA. Introduction of self-healing moieties not only improved the residence time and lubrication properties of HA, but also endowed the precursors with biological functions otherwise lacking. In particular, self-healing HA showed improved viscoelastic properties without compromising its injectability. These modified HA molecules also exhibited enhanced free radical scavenging and diminished enzymatic degradation. Moreover, intraarticular administration of self-healing HA can be used to slow the progression of cartilage degeneration after trauma. In this proof-of-concept study, we used low molecular weight (200 kDa) HA as the precursor, which was necessary to limit the confounding contributions arising from high molecular weight HA. Having demonstrated the promising beneficial effect of self-healing HA, future studies will include self-healing HA generated from high molecular weight HA precursors in a larger animal model. Our approach of using molecular engineering to imbue biomaterials with self-healing could serve as a guide for engineering self-regenerating biomaterials and lubricants with greater efficacy and therapeutic outcome.

## Methods

### Synthesis of UPy-bearing linker

UPy-bearing linkers were synthesized *via* a multi-step process.^[11a, 30]^ Briefly, 2-amino-4-hydroxy-6-methylpyrimidine (Sigma, Cat.# A58003) (10 g, 0.08 mol) was dissolved in excess 1,6-diisocyanatohexane (Sigma, Cat.# 52650) (107 g, 0.64 mol) and reacted at 100°C overnight in argon environment. The product, termed as compound **1** — 1-(6-isocyanatohexyl)-3-(6-methyl-4-oxo-1,4-dihydropyrimidin-2-yl) urea in Fig. 1b, was precipitated in n-pentane, filtered, and dried. The product was characterized by using NMR and FTIR (**Supplementary Figs. 1 and 5**). Compound **1** (5 g, 0.017 mol) was mixed with *N*-boc-1,6-hexanediamine (TCI Chemicals, Cat.# A1375) (5.5 g, 0.025 mol) in anhydrous dichloromethane (∼75 mL) and kept at 50°C overnight to yield compound **2** (tert-butyl(6-(3-(6-(3-(6-methyl-4-oxo-1,4-dihydropyrimidin-2-yl)ureido)hexyl) ureido)hexyl)carbamate), which was precipitated in chilled diethyl ether, filtered, and dried. NMR and FTIR spectra are provided in supplementary information (**Supplementary Figs. 2 and 5**). Compound **2** (5 g) was dispersed in dichloromethane (90 mL). Trifluoroacetic acid (TCI Chemicals, Cat.# A12198) (10 mL) was added to the suspension and stirred vigorously at room temperature for ∼6 h. Following the reaction, the dichloromethane and trifluoroacetic acid were removed using a rotavapor. The solid residue was dissolved in minimum amount of dichloromethane and precipitated in excess chilled acetone. The product, compound **3**, (1-(6-(3-(6-aminohexyl)ureido)hexyl)-3-(6-methyl-4-oxo-1,4-dihydropyrimidin-2-yl)urea·trifluoroacetic acid) was filtered, washed repeatedly with acetone and dried in vacuum. The product was characterized by NMR and FTIR analyses (**Supplementary Figs. 3 and 5**). The dried compound **3** was treated with Amberlite IRA 400 chloride ion exchange resin (Sigma, Cat.# 247669) in dimethylsulfoxide-water mixture (1:1) at room temperature for 2 h. The resin was filtered off and the solution of the UPy-bearing linker (compound **4**, 1-(6-(3-(6-aminohexyl)ureido)hexyl)-3-(6-methyl-4-oxo-1,4-dihydropyrimidin -2-yl)urea·HCl) was used to react with HA. The product was characterized by NMR and FTIR spectra (**Supplementary Figs. 4 and 5**).

### Synthesis of HA-UPy

To synthesize HA-UPy, UPy-bearing linker was reacted with sodium hyaluronate (HA, MW 200 kDa or 1MDa; Lifecore Biomedical, Cat# HA200K) using EDC/NHS chemistry. Briefly, HA was dissolved in a mixture of deionized water and DMSO (1:1) at 5 mg/mL, to which 1-ethyl-3-(3-dimethylaminopropyl)carbodiimide hydrochloride (EDC, TCI Chemicals, Cat.# D1601), *N*-hydroxysuccinimide (NHS, Sigma, Cat.# 130672), and compound **4** (each 1 equivalent with respect to the carboxylic acid groups of HA) were sequentially added at 15 min intervals. The reaction was carried out at room temperature for 48 h, and the resulting HA-UPy product was purified *via* dialysis against water and lyophilized. Successful conjugation of UPy to HA was verified by a combination of FTIR and NMR spectroscopy, and the extent of UPy conjugation was quantified *via* ^1^HNMR spectroscopy.

### Synthesis of HA-Cy 7 and HA-UPy-Cy7

Cyanine 7 (Cy7, Lumiprobe, Cat.# 550C0) dye was conjugated onto HA and HA-UPy *via* amide coupling reaction. Briefly, HA or HA-UPy was dissolved in a mixture of deionized water and DMSO (1:1) at 5 mg/mL. EDC (1 equivalent with respect to the carboxylic acid group of HA or HA-UPy), NHS (1 equivalent with respect to the carboxylic acid group of HA or HA-UPy), and Cy7 (0.04 equivalent with respect to the carboxylic acid group of HA or HA-UPy) were subsequently added to the polymer solution. After 48 h of reaction at room temperature, the mixture was dialyzed extensively with water for 4 d. The solutions were then freeze-dried to obtain HA-Cy7 or HA-UPy-Cy7. The product was characterized by a combination of FTIR and NMR spectroscopy, and the extent of dye conjugation was quantified *via* UV/vis absorption spectroscopy.

### Fourier transform infrared (FTIR) and nuclear magnetic resonance (^1^HNMR) spectroscopy>

FTIR spectrometer (Nicolet 8700) with an attenuated total reflection (ATR) range of 4000 to 650 cm^-1^ was used for all characterization. For ^1^HNMR measurements, the products were dissolved in heavy water (∼1 wt%), and the spectra were recorded by using a 500 MHz Agilent/Varian VNMRS spectrometer at either room temperature or 80°C.

### Gelation of HA and HA-UPy molecules

HA and HA-UPy solutions of various concentrations (2 wt%, 5 wt%, and 10 wt%) were prepared by dissolving the required weight of the molecules in phosphate-buffered saline (PBS). For visualizing gelation, food dye was added. Eppendorf tubes containing the solutions were inverted to visualize the flow under gravity, and images were taken both immediately following dissolution and after 24 h.

### Rheological analysis

Both HA-UPy and HA were prepared in PBS and subjected to rheological measurements as a function of concentration by using a rotational rheometer (AR-G2, TA Instruments). Each sample was loaded on a parallel plate geometry (Al, diameter 8 mm), and the oscillatory frequency sweep measurements were conducted at 1% strain amplitude with frequencies ranging from 0.1 to 10 Hz. To assess the shear-thinning behavior, the steady-state viscosities of HA-UPy and HA at 10 wt% were measured at 1% strain as a function of shear rate (0.1 to 100 s^-1^). To evaluate the recovery of HA-UPy and HA at 10 wt%, step-strain measurements were recorded at 1 rad/s with a range of consecutive strains (1%, 100%, 1%, 1000%, and 1%) applied each for 180 s. To examine hysteresis of HA-UPy, 6 cycles of alternative low (1%) and high (500%) strain were applied. All samples were measured in triplicate.

### Injectability of HA-UPy molecules

10 wt% HA-UPy molecules were generated in PBS, loaded into a Hamilton syringe, and extruded into different shapes through a 26G needle.

### Self-healing of HA-UPy molecules

To examine the self-healing phenomenon, hydrogel pieces were generated from HA-UPy (10 wt% and colored differently for visualization). Several pieces of the hydrogel were gently brought into contact with one another.

### Enzymatic degradation

The stability of HA-UPy was evaluated in the presence of hyaluronidase (Sigma, Cat.# H3506).^[21]^ In brief, HA-UPy and HA were dissolved in 20 mM sodium acetate buffered solution (pH 6) at 2.5 mg/mL supplemented with 1 kU/mL hyaluronidase. Each sample was sealed in a benzoylated dialysis membrane (MWCO ∼ 2 kDa; Sigma, Cat.# D2272) and dialyzed against sodium acetate buffer at 37°C for 48 h. The dialysate containing the degradation products was collected for uronic acid assay. The solution was mixed with 12.5 mM sodium tetraborate (Alfa Aesar, Cat.# A16176) in concentrated sulfuric acid at a volume ratio of 1:6 and heated at 100°C for 10 mins. Upon cooling, 0.15% m-hydroxydiphenyl (Sigma, Cat.# 262250) in 0.5% NaOH was added and its absorbance was measured at 520 nm using UV/vis spectroscopy. Known concentrations of HA (0 − 2.5 mg/mL) were used to generate the standard curve.

### Free radical scavenging

The ability of HA-UPy to scavenge free radicals was analyzed by using 1,1-diphenyl-2-picrylhydrazyl (DPPH; Sigma, Cat.# D9132) or Fenton reagent as free radical sources.^[16]^ For the DPPH assay, 10 wt% HA-UPy or HA was fully soaked in ethanol containing 0.1 mM DPPH at 37°C for 1 h in the dark. Saline of the same volume was used as the control. The absorbance of DPPH solution at 517 nm before (Abs_0_) and after (Abs_t_) the incubation was recorded using UV/vis. The DPPH scavenging effect was determined as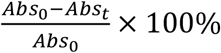. The average reduction in absorbance in the saline group was subtracted from the HA and HA-UPy groups to normalize for dilution. All samples were measured in triplicate.

To examine the hydroxyl radical scavenging effect, the Fenton reagent was prepared as described elsewhere.^[31]^ Briefly, a reaction mixture consisting of 1 mM ferric chloride (Sigma, Cat.# 157740), 30 mM deoxyribose (Sigma, Cat.# 121649), 1 mM ascorbic acid (Sigma, Cat.# A92902), 1 mM EDTA (Sigma, Cat.# E9884) and 20 mM H_2_O_2_ was prepared in 0.2 M phosphate buffer. 1 mL of the reaction mixture was added to either 100 µL of 10 wt% HA-UPy or 0.2 M phosphate buffer (control). The gel or phosphate buffer were incubated at room temperature for 1 h on a shaker plate. Following the incubation period, 500 µL of the reaction mixture was taken from each tube and mixed with 500 µL of a solution of 0.25% thiobarbituric acid (TBA; Sigma, Cat.# T5500) in 15% trichloroacetic acid (TCA; Sigma, Cat.# T6399). The mixtures were incubated in a silicon oil bath at 100°C for 20 min. The absorbance was measured at 532 nm using a UV/Vis spectrophotometer, with a lower absorbance corresponding to fewer hydroxyl radicals. All samples were measured in triplicate.

Following exposure to the Fenton reaction mixture, the samples were washed in PBS, freeze dried, and reconstituted in saline at a concentration of 10 wt%. Corresponding HA-UPy samples were similarly incubated in phosphate buffer alone (controls), followed by washing in PBS, freeze drying, and reconstituting in saline. A frequency sweep was performed as previously described at 1% strain under oscillatory mode with frequency varying from 0.1 to 10 Hz. The storage moduli (G′) at 1 Hz was compared for the free radical-treated HA-UPy and corresponding HA-UPy control.

### Explant coculture

To examine the biocompatibility of HA-UPy, rat tibial condyles were harvested and cultured in chondrocyte medium with or without HA-UPy (10 wt%, 50 μL) for 7 d. The cartilage explants were then rinsed and incubated in PBS containing 0.05% calcein acetoxymethyl and 0.2% ethidium homodimer-1 from the Live/Dead Cell Viability Assays kit (Life technologies, Cat.# L3224) for 30 min. After thorough washing, the explants were sectioned and imaged using a Keyence (BZ-X710) microscope.

### Coefficient of friction between cartilage explants

Two flat cartilage discs (8-mm diameter and 0.5-mm thickness, porcine, 3-year-old) were mounted on sandpaper-covered parallel plates using cyanoacrylate glue, with the articular surfaces facing each other. After equilibration in saline, HA, HA-UPy (10 wt%), or saline was introduced at the cartilage-cartilage interface, the discs were brought into contact and programmed to move against each other by using a rotational rheometer. Both normal stress and shear stress were recorded under shear rates ranging from 0.1 to 1 s^-1^. The coefficient of friction (μ) at the cartilage-cartilage interface was calculated using the equation,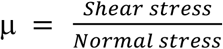.

### ACL injury models

All animal studies were approved by the Institutional Animal Care and Use Committee at Duke University in compliance with NIH guidelines for laboratory animal care. Both female mice (C57BL/6J, 3-month-old, Jackson Lab) and rats (Lewis, 3-month-old, Charles River) were used for unilateral anterior cruciate ligament transection (ACLT). Each animal was sedated using 2% isoflurane and injected with buprenorphine (1 mg/kg, sustained release, ZooPharm) as an analgesic prior to surgery.

For ACLT in mouse, each animal was placed in a supine position with the left hindlimb bent over a triangular cradle. After shaving and disinfecting the skin, a cut less than 0.5-cm-long was created from the medial side to expose the joint capsule. The ACL was fully extended by bending the knee to 90°C and transected using spring scissors (FST, Cat.# 15004-08). Bupivacaine (0.5%, Hospira) was then applied topically, and the incision was closed with Vicryl 5-0 sutures.

For mi-ACLT in rat, the left hindlimb was disinfected and flexed to approximately a 90° angle. An 18G needle was inserted perpendicularly into the joint on the lateral side of the patellar ligament, and the bevel of the needle was used to transect the ACL. To confirm the completion of ACLT, the clinical anterior drawer test was performed. In the anterior drawer test, the tibia easily moved out of the normal range of motion when pressure was applied behind the tibia while the femur was held in place.

Following surgery, animals were placed on heating pads to aid in recovery from anesthesia and left unconstrained in cages.

### Intra-articular injections

Sterile saline, HA (200 kDa, 10 wt% in saline), and HA-UPy (200 kDa, 10 wt% in saline) were prepared fresh and injected into the injured joints using a Hamilton syringe fitted with 26G needle. Animals were randomized to treatment groups following surgery. For mouse, injections were started one week following ACLT and performed weekly for four weeks with 5 μL of sample injected each time. For rat, injections were started one day following mi-ACLT and repeated weekly for another eight weeks with 50 μL of sample injected each time. One week after the last injection, animals were euthanized.

### IVIS imaging

Calibration studies with varying extent of Cy7 modified HA molecules were carried out to identify optimal Cy7 concentration required to obtain the optical intensity. 50 μL of Cy7-containing HA or HA-UPy (10 wt%) was administered into the rat joint via intra-articular injection. At designated times after injection, rats were anesthetized under isoflurane inhalation and imaged by using an IVIS Kinetics system (excitation filter, 745 nm; emission filter, ICG; excitation time, 100 ms). The epi-fluorescence intensity of Cy7 in the joint was quantified by selecting an ROI from images taken at Day 0. The percent of initial fluorescence intensity was calculated after measuring the fluorescence intensity at each subsequent time point.

### Histological analysis

Fixed joint samples were decalcified in 14% EDTA (pH 8.0, Sigma, Cat.# E5134) for 2–3 weeks, rinsed with PBS, dehydrated, and embedded in paraffin blocks, and sliced into 8 μm-thick sections using a Leica rotary microtome. Each section was deparaffinized in CitriSolv (Decon Labs, Cat.# 1601) and rehydrated through graded alcohols and deionized water. For Safranin-O staining, the rehydrated sections were incubated in 1% Safranin-O (Sigma, Cat.# S8884) for 30 min for mouse cartilage sections and 1 h for rat cartilage sections. They were counter-stained with 0.02% Fast Green (Sigma, Cat.# F7258) for 1 min for mouse and 1.5 min for rat, followed by incubating in hematoxylin solution (Ricca Chemical, Cat.# 3536-16) for 1 min for mouse and 5 min for rat. The stained sections were rinsed, dehydrated, and covered with a mounting medium (Fisher Scientific, Cat.# SP15-100). Images were taken using a Keyence microscope.

### OARSI scoring

Scoring of Safranin O-stained sagittal sections was performed by three blinded scorers in accordance with the OARSI histopathology initiative for evaluation of OA in mouse^[32]^ and rat.^[25]^ Scoring criteria for the mouse included the degree of degeneration of cartilage in both the tibia and femur, evaluated from 0-6: 0 means normal; 0.5 has loss of proteoglycan without structural changes; 1 shows limited fibrillation on the cartilage surface; 2 presents vertical clefts; 3 means vertical clefts or erosion covering <25% of surface area; 4 for 25–50% area being affected; 5 for 50–75%; and 6 for >75%. Scoring criteria for the rat included total tibial cartilage degeneration (0-5 for 3 zones, total 0-15), femoral cartilage degeneration (0-5), bone score (0-5), and osteophyte score (0-4), with total added score from 0-29. The osteophyte score was modified for sagittal joint sections based upon a histogram of osteophyte sizes for all samples in the mi-ACLT groups. ImageJ was used to quantify total cartilage degeneration, significant cartilage degeneration, and surface matrix loss (all expressed as the percentage of total cartilage width), as well as depth ratio of cartilage lesions (expressed as the depth of degenerated cartilage to the thickness of total cartilage). These measurements were completed for both the tibia and femur, the latter of which is a modification of the original scoring which only examines the tibia. A total of four sections, taken from two locations within the medial compartment of the joint, were evaluated. These four scores/measurements were averaged as technical replicates for each subject.

### Immunohistochemical analysis

Immunohistochemical staining was used to detect MMP-13 and ADAMTS-5 expression in cartilage. Briefly, deparaffinized sections were subjected to heat-induced antigen retrieval in a vegetable steamer for 13 min and permeabilized with 0.1% Triton X-100 for 15 min. Tissue sections were blocked with 0.1% BSA and 0.26 M glycerol in TBS for 1 h, sequentially exposed to Dual Endogenous Enzyme Block (Dako, Cat.# S2003) for 30 min, and 5% and 1.5% normal goat serum for 30 min and 1 h, respectively, before incubation at 4°C overnight with either anti-MMP13 (dilution 1:500, Abcam, Cat.# ab39012) or anti-ADAMTS5 (dilution 1:100, Abcam, Cat.# ab41037). VECTASTAIN^®^ Elite ABC HRP Kit (Vector Laboratories, Cat.# PK-6100) kit and ImmPACT® DAB Peroxidase Substrate (Vector Laboratories, Cat.# SK-4105) were used to visualize the enzyme expression.

### Statistical analysis

The means with standard error of mean (n ≥ 3) are presented in the results. All animal studies included at least 6 animals per group. All the data were subjected to two-tailed Student’s t-test, one-way analysis of variance (ANOVA) with post-hoc Tukey’s multiple comparisons test, or two-way repeated measures ANOVA with Tukey’s multiple comparisons test using GraphPad Prism 8. Any p-value of less than 0.05 was indicated with an asterisk and was considered statistically significant. All experiments were reproduced independently.

## Supplementary Information

Supplementary Figures 1-12

Supplementary Text

Supplementary Methods

## Acknowledgments

We thank H Newman, S Sharma, and N Sangaj for helping the with the blinded OARSI scoring. This material is based upon work partially supported by the National Science Foundation Graduate Research Fellowship Program (A Gilpin) under Grant No. DGE 1644868. This work was partially supported by the Duke University and National Institute of Arthritis and Musculoskeletal and Skin Diseases (NIAMS) of the National Institutes of Health (NIH) under Award Number NIH R01 AR063184. The content is solely the responsibility of the authors and does not necessarily represent the official views of the funding agencies.

## Author contributions

S.V conceived the initial idea. A.G., Y.Z., and S.V. designed the study and wrote the manuscript. A.G., Y.Z., and J.H. designed and performed experiments and analyzed the data. J.H.R performed/ helped with synthesis. Y.Y and Y.Z performed the mouse studies. W.E performed/helped with the rat ACL injuries. S.Z helped with the rheological measurements. All authors were involved in the manuscript preparation

