## Supplemental Data for "Self-healing of hyaluronic acid to improve *in vivo* retention and function"

#### Supplementary Information

Supplementary Figures 1-12

Supplementary Note

Supplementary Methods

#### Supplementary Figures

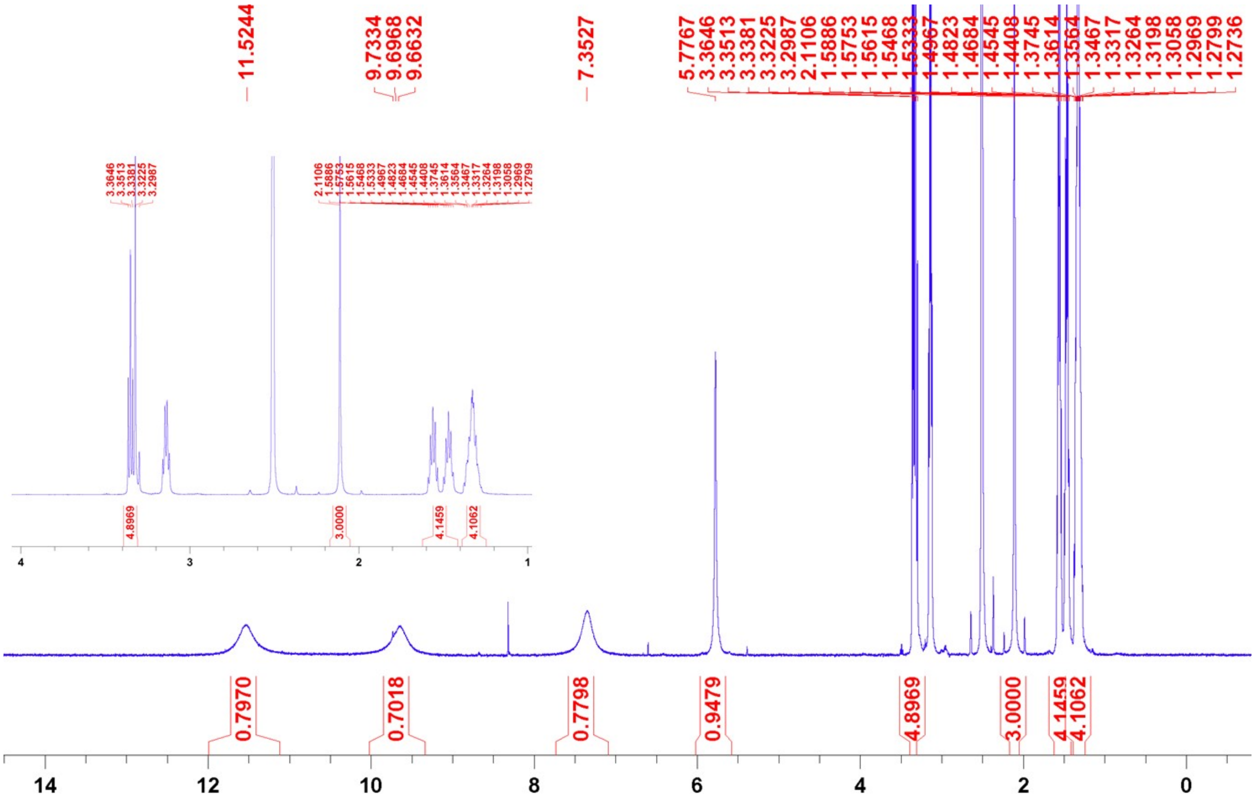

**Supplementary Figure 1.** $^{1}$HNMR Spectrum of compound 1 recorded in DMSO-$d_6$ at room temperature. Extended spectrum from 1-4 ppm (Inset).

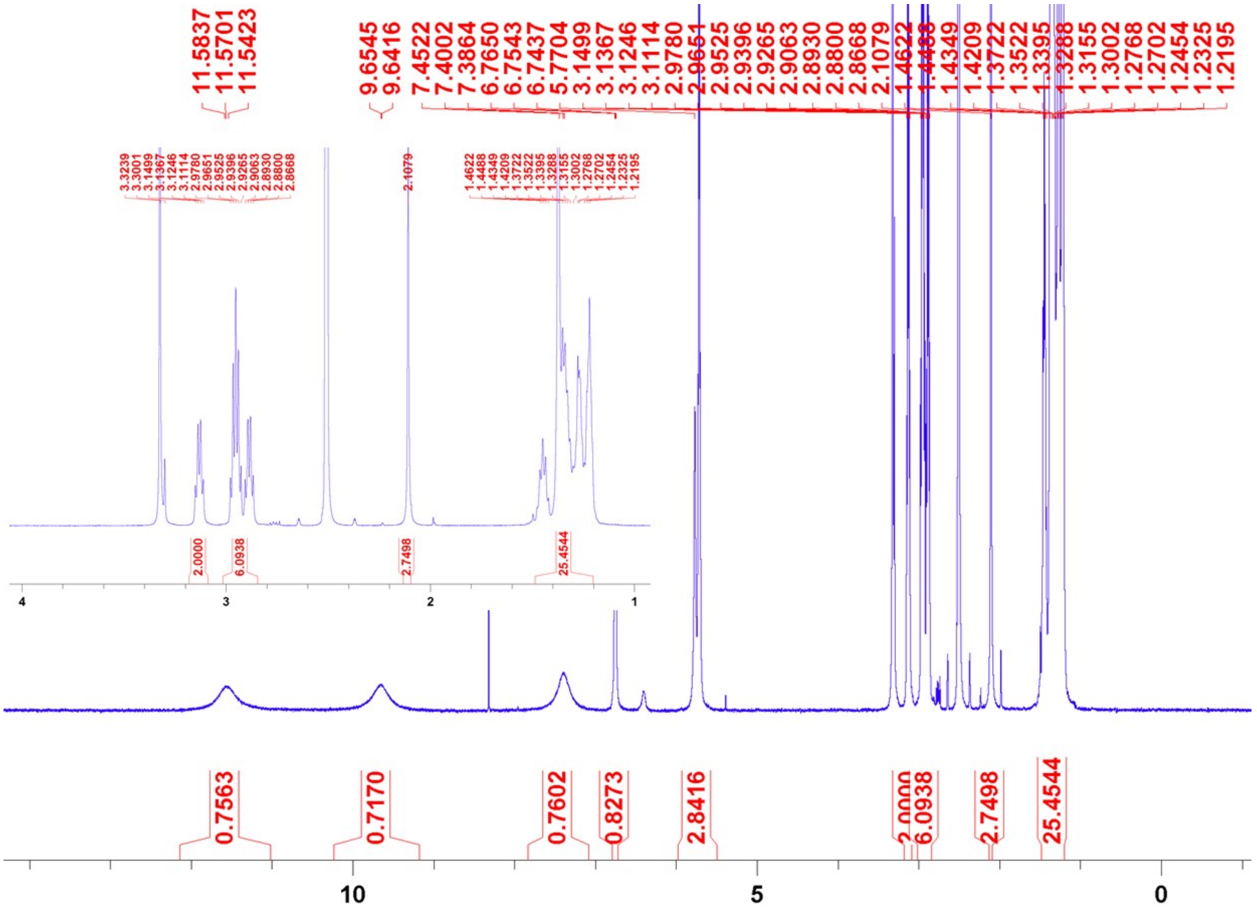


**Supplementary Figure 2.** $^{1}$HNMR Spectrum of compound 2 recorded in DMSO-$d_6$ at room temperature. Extended spectrum from 1-4 ppm (Inset).

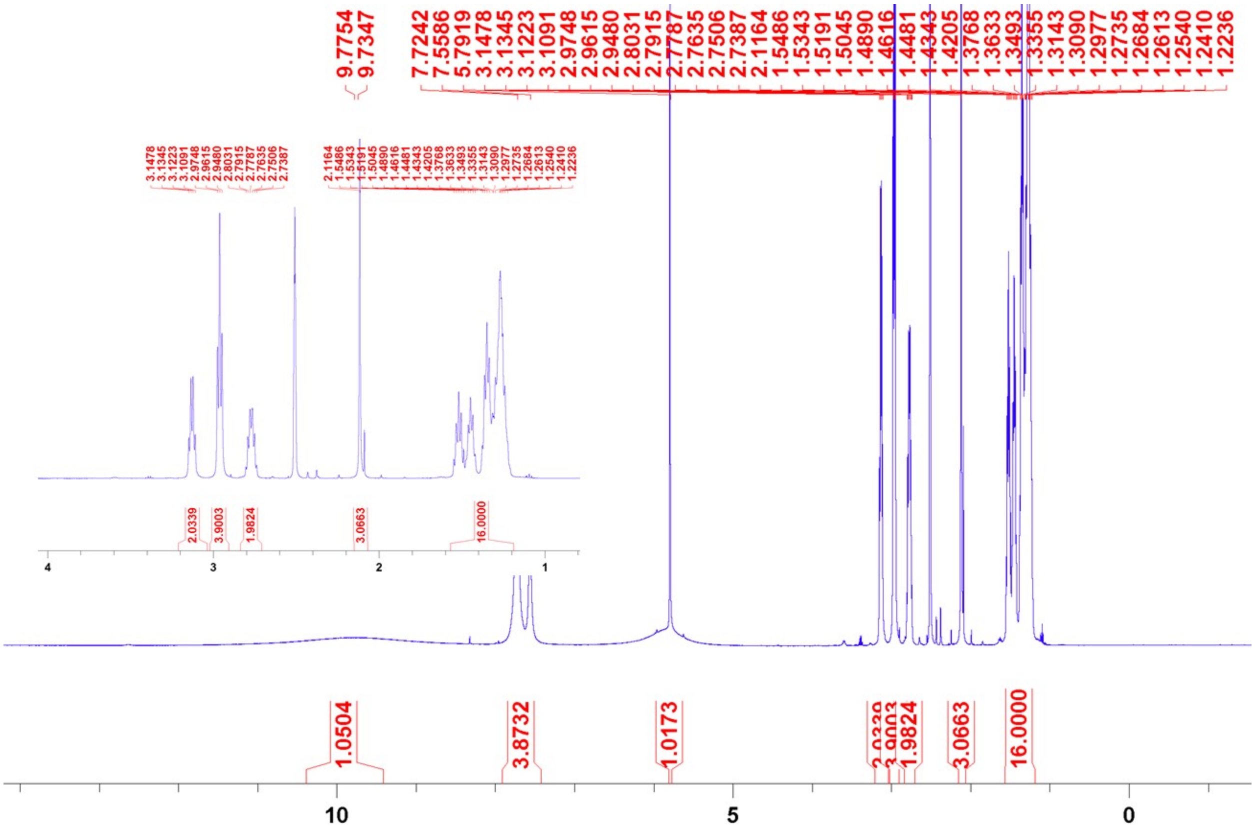


**Supplementary Figure 3.** $^{1}$HNMR Spectrum of compound 3 recorded in DMSO-$d_6$ at room temperature. Extended spectrum from 1-4 ppm (Inset).

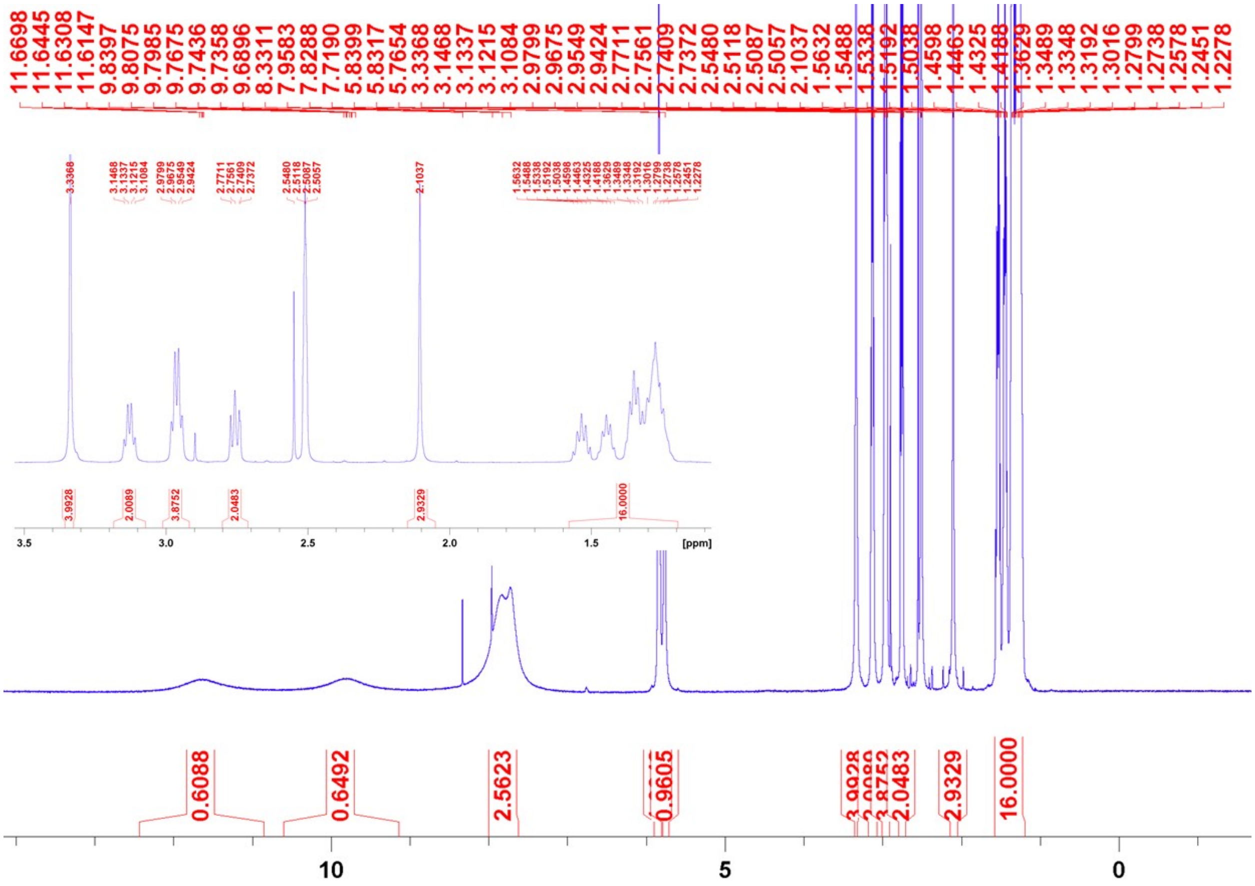


**Supplementary Figure 4.** $^{1}$HNMR Spectrum of compound 4 recorded in DMSO-$d_6$ at room temperature. Extended spectrum from 1.5-3.5 ppm (Inset).

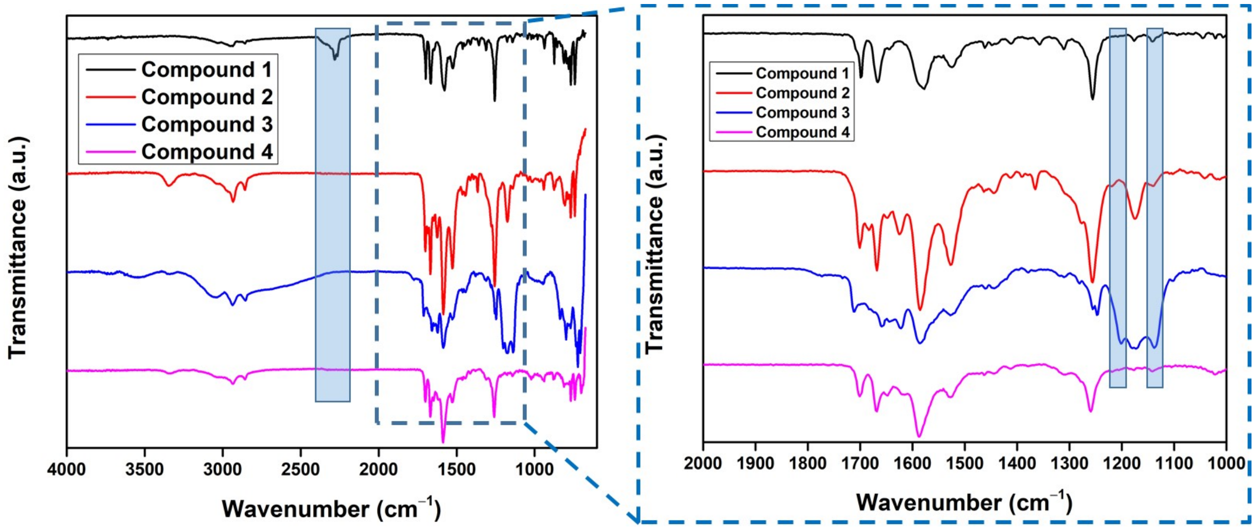


**Supplementary Figure 5.** FTIR (ATR) spectra of compound 1-4 recorded at room temperature. Left panel: full spectrum from 4000-600 $cm^{-1}$. The spectra clearly show the presence of NCO stretching frequency for compound 1 at 2271 $cm^{-1}$ (marked with transparent color box). Right panel shows the extended spectrum from 2000-1000 $cm^{-1}$. The spectra show the presence of UPy stretching frequencies at 1699, 1669, 1575 and 1520 $cm^{-1}$ respectively for all the compounds. In addition, compound 3 shows the presence of bands at 1201 and 1135 $cm^{-1}$ respectively corresponding to TFA absorptions (marked with transparent color boxes).

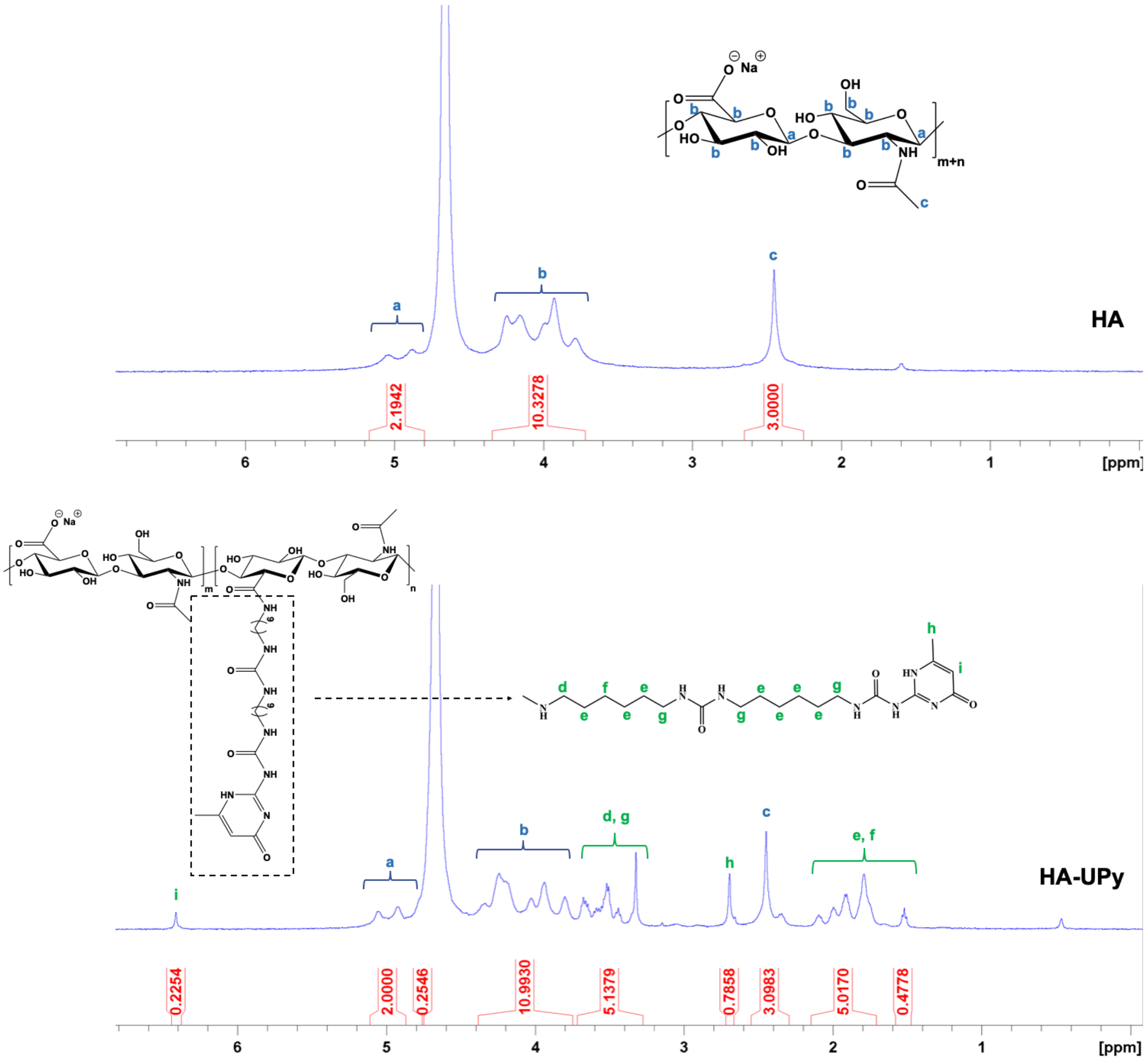


**Supplementary Figure 6. $^1$H NMR analysis of HA and HA-UPy.**

The protons were labeled alphabetically and assigned to the peaks, the integrated areas of which were normalized. To calculate the grafting density (GD) of UPy, the signature peaks of UPy at δ 6.41 ppm (i, 1H) and δ 2.69 ppm (h, 3H) from the pyrimidinone protons were compared against the signature peaks of HA at δ 4.92-5.06 ppm (a, 2H) and δ 2.45 ppm (c, 3H) from the anomeric carbon-attached and acetyl protons. $GD_1 = \frac{i/1}{c/3} = \frac{0.2254/1}{3.0983/3} \times 100\% = 21.8\%$ of HA's dimeric repeating units. $GD_2 = \frac{h/3}{c/3} = \frac{0.7858/3}{3.0983/3} \times 100\% = 25.4\%$ of HA's dimeric repeating units.

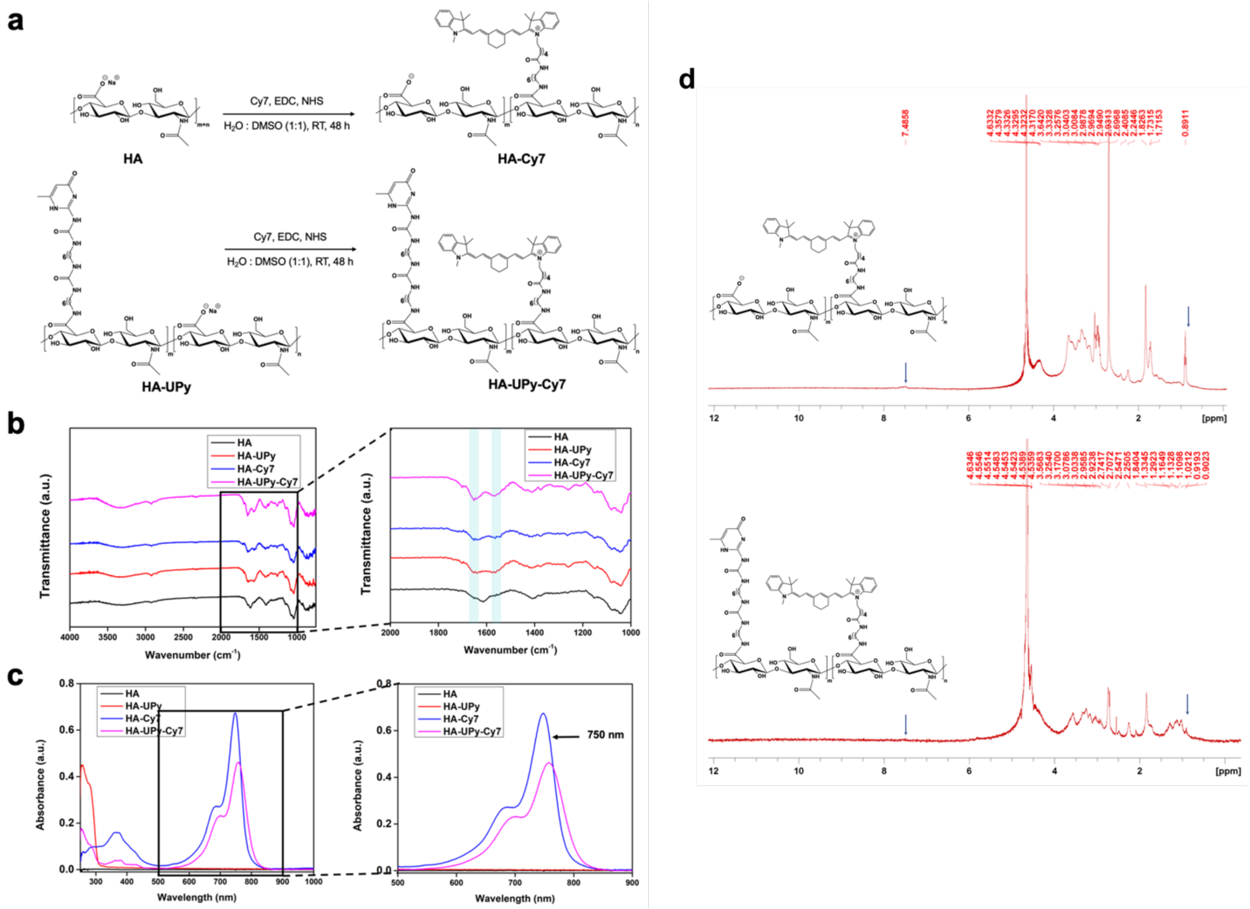

**Supplementary Figure 7. Synthesis and characterization of Cy7-tagged HA and HA-UPy.**

***a,*** Synthesis scheme of HA-Cy7 and HA-UPy-Cy7. ***b,*** FTIR spectra of Cy7-tagged polymers show absorptions at 1656 $cm^{-1}$ and 1564 $cm^{-1}$ (shaded bands), characteristic of amide I and amide II stretching from Cy7, as opposed to a broad absorption peak at 1612 $cm^{-1}$ that corresponds to C=O stretching of amide group from HA. ***c,*** The extent of dye conjugation was calculated using UV/vis absorption spectroscopy with a standard calibration curve of Cy7 absorbance at 750 nm. The dye content was found to be 4.5 wt% for HA-Cy7 and 3.9 wt% for HA-UPy-Cy7. ***d,*** $^{1}H$ NMR spectroscopy of HA-Cy7 and HA-UPy-Cy7 reveals the presence of aromatic protons at δ 7.48 ppm from the benzene ring and methyl proton at ~0.9 ppm of Cy7.

a

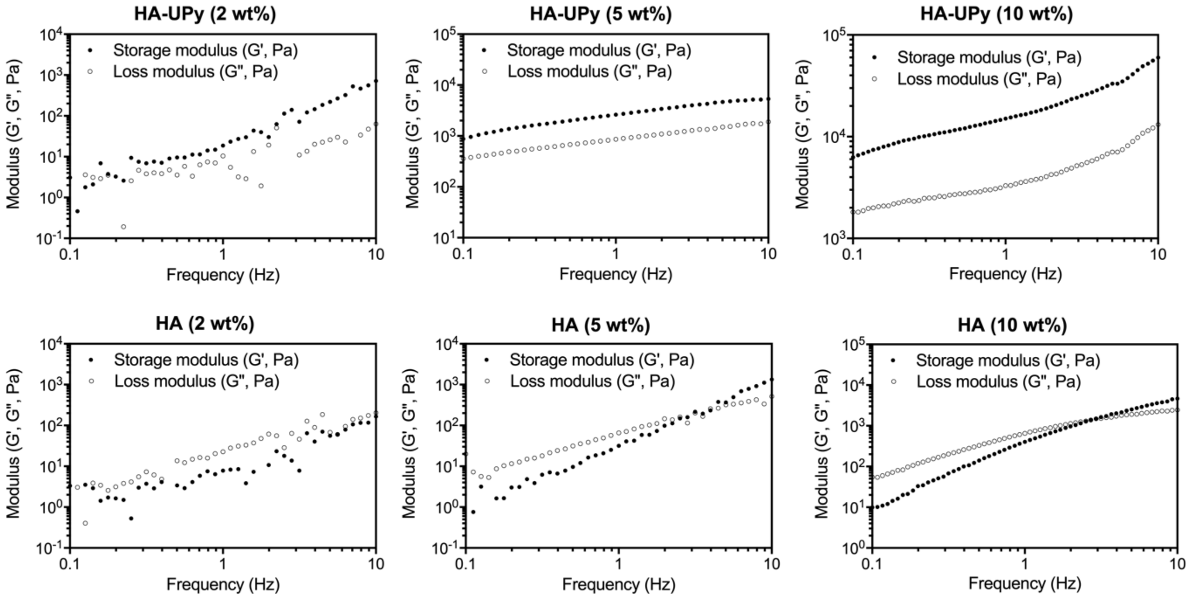


b

| Concentration | Sample | G' (Pa) | G" (Pa) | tan (delta) |
|---|---|---|---|---|
| 2 wt% | HA | 12 ± 9 | 16 ± 5 | N/A |
| | HA-UPy | 49 ± 20 | 27 ± 10 | 0.58 ± 0.05 |
| 5 wt% | HA | 51 ± 30 | 95 ± 50 | 1.9 ± 0.3 |
| | HA-UPy | 1500 ± 600 | 550 ± 200 | 0.38 ± 0.03 |
| 10 wt% | HA | 260 ± 80 | 470 ± 100 | 2.0 ± 0.2 |
| | HA-UPy | 16000 ± 3000 | 3500 ± 700 | 0.21 ± 0.004 |

c

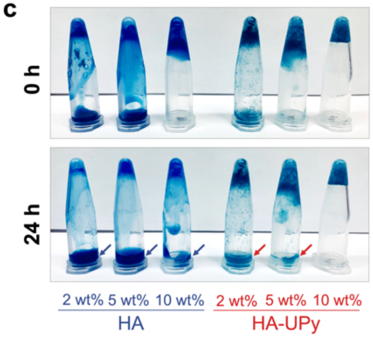


**Supplementary Figure 8. Rheological properties of HA and HA-UPy.**

***a,*** Frequency sweep measurements for HA and HA-UPy at different concentrations show the evolution of storage (G′) and loss (G″) moduli as a function of frequency. ***b,*** G′, G″, and delta of HA and HA-UPy at different concentrations measured at 1 Hz. ***c,*** Photos of HA-UPy and HA as a function of concentration.

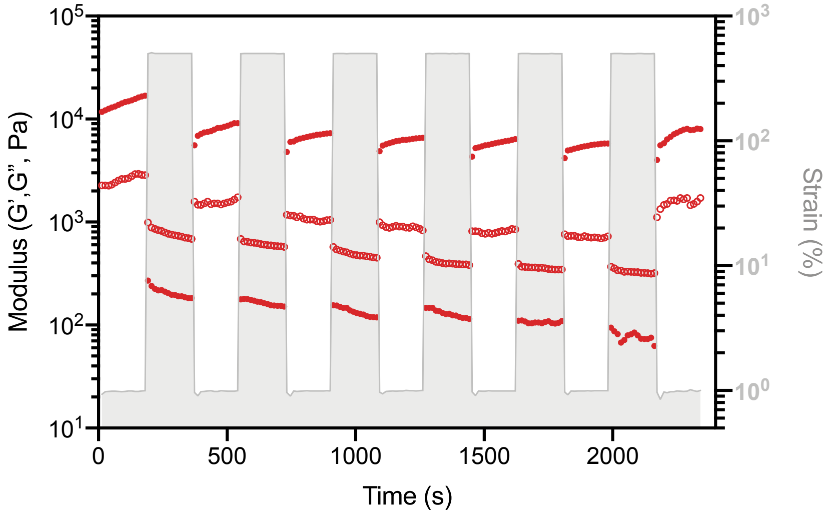


**Supplementary Figure 9. Recovery of HA-UPy network.**

HA-UPy returns to original G′ value following six cycles of low and high strains, indicating complete recovery of the network.

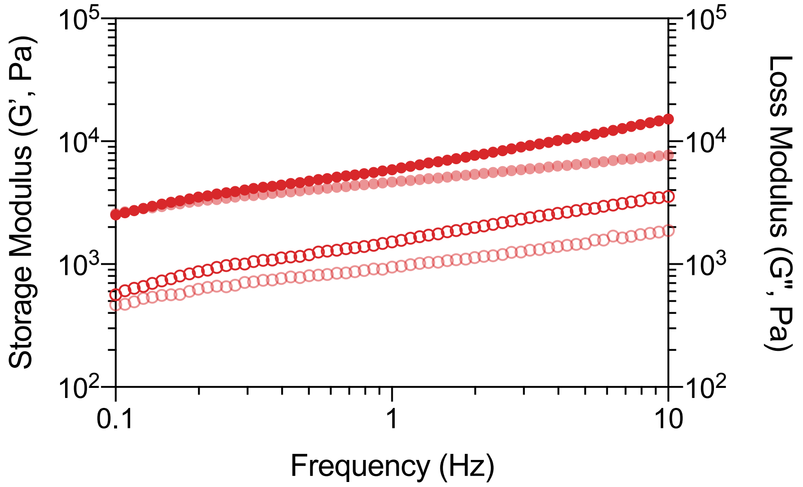


**Supplementary Figure 10. Frequency sweep of free radical-exposed HA-UPy.**

Storage (G′) and loss (G") modulus of HA-UPy following exposure to hydroxyl radicals as a function of frequency. Minimal reduction in G′ and G′′ are observed compared to control HA-UPy.

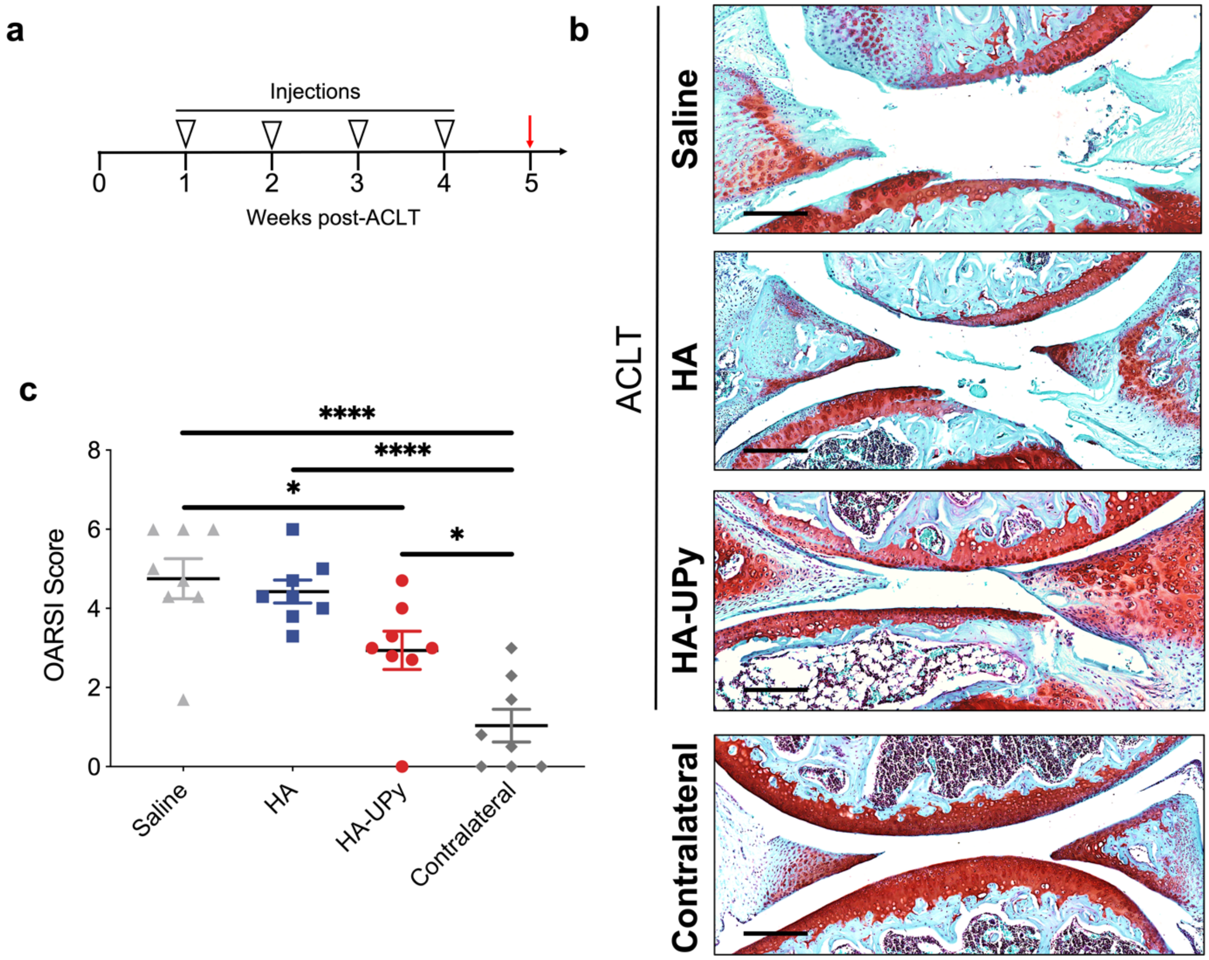


**Supplementary Figure 11. Chondroprotection of HA-UPy in mouse surgical ACLT model.** ***a,*** Experimental timeline showing schedule of injections of saline, HA, or HA-UPy. ***b,*** Morphology change of mouse joints at week 5 post-ACLT. Contralateral joints without injury were used as positive control. Scale bars: 200 μm ***c,*** OARSI scores of mouse joints based on the Safranin-O staining results. Data are presented as means (± s.e.m.). One-way ANOVA with Tukey's multiple-comparisons test was used for statistical analysis. Significance is determined as *$P < 0.05$, ****$P < 0.0001$.

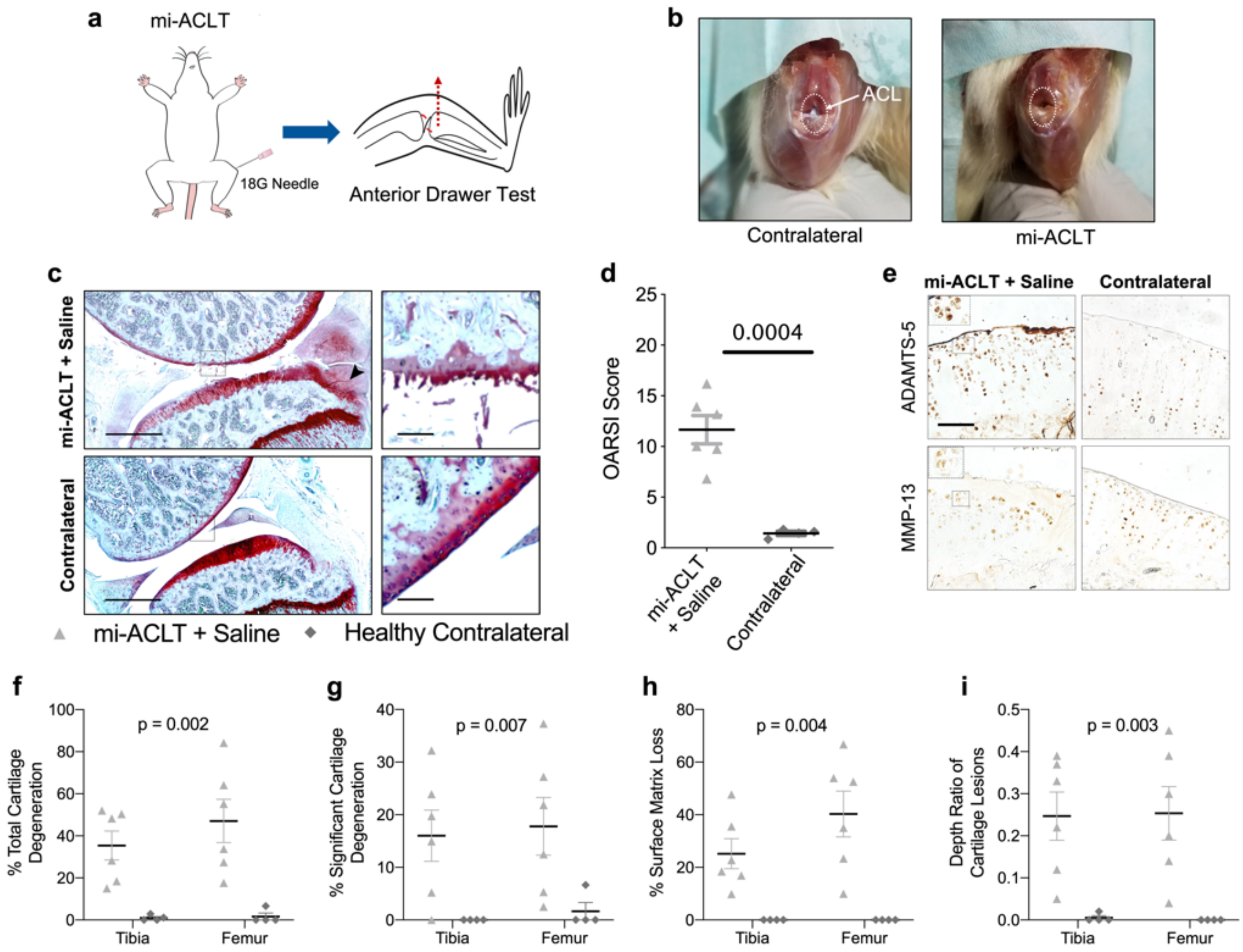


**Supplementary Figure 12. Minimally invasive ACLT (mi-ACLT).**

***a,*** The rat mi-ACLT procedure was performed on the left hind knee joint while the knee was flexed. ACLT was confirmed using anterior drawer test, whereby the tibia protrudes from the joint when ruptured. ***b,*** Dissected knee joints with intact ACL and transected ACL. ***c,*** Safranin O-stained knee joint from mi-ACLT group following 8 weeks of sham saline injections show severe cartilage degeneration and an osteophyte of the tibia (arrow). Contralateral joints without injury were used as positive control. Scale bar: 1 mm. ***e,*** The unoperated contralateral joint has a smooth cartilage surface and no osteophytes. ***d,*** OARSI scoring of the knee joints shows that the mi-ACLT group has a significantly higher degree of cartilage degeneration than the contralateral group. Data are presented as means (± s.e.m.). A two-tailed (unpaired) t-test was used to analyze statistical significance between groups. ***e,*** The cartilage of the mi-ACLT group shows more cells positive for ADAMTS-5 and MMP-13 expression, as well as clustering of chondrocytes (inset). ***f-i,*** Quantitative measures of ***(f)*** total cartilage degeneration, ***(g)*** significant cartilage degeneration, ***(h)*** cartilage surface matrix loss, and ***(i)*** thickness of cartilage lesions were greater in the mi-ACLT group than the contralateral measures for both the tibia and femur. Data are presented as means (± s.e.m.). A two-way repeated measures ANOVA was used to analyze statistical significance. The significance of treatment effect is shown above the graphs.

### Supplementary Note

**<u>Characterization of UPy linker and intermediate products:</u>**

As shown in Fig. 1b, the synthesis of UPy linker (compound 4 in Fig. 1b) was a multistep process involving three intermediate compounds (Compounds 1-3). Each compound was characterized by $^1$HNMR and FTIR following the reaction. $^1$HNMR was carried out by dissolving each compound in DMSO-d$_6$, and the spectra were recorded by using a 500 MHz Agilent/Varian VNMRS spectrometer at room temperature.

***Characterization data for compound 1:***

$^1$HNMR (DMSO-d$_6$): δ (ppm) = 11.52 (s, 1H, –(CH$_3$)C–*NH*–), 9.69 (s, 1H, –CH$_2$–NH(CO)–*NH*–), 7.35 (s, 1H, –CH$_2$–*NH*(CO)–NH–), 5.77 (s, 1H, –*CH*=C(CH$_3$)–), 3.29-3.36 (m, 4H, –NH–(CO)–NH–*CH$_2$*– + –*CH$_2$*–NCO), 2.11 (s, 3H, –CH$_3$), 1.44-158 (m, 4H, –CH$_2$–*CH$_2$*–(CH$_2$)$_2$–*CH$_2$*–CH$_2$–NCO), 1.27-1.37 (m, 4H, –CH$_2$–CH$_2$–*(CH$_2$)$_2$*–CH$_2$–CH$_2$–NCO).

FTIR (ATR): ν (cm$^{-1}$) = 2271 (NCO stretch), 1696 (UPy), 1666 (UPy), 1576 (UPy), 1522 (UPy), 1464, 1357, 1312, 1252.

***Characterization data for compound 2:***

$^1$HNMR (DMSO-d$_6$): δ (ppm) = 11.57 (br.s, 1H, –(CH$_3$)C–*NH*–), 9.65 (s, 1H, –CH$_2$–NH(CO)–*NH*–), 7.40 (br. s, 1H, –CH$_2$–*NH*(CO)–NH–), 6.75 (m, 3H, –C(O)–*NH*(CH$_2$)$_6$*NH*–C(O)– + (CH$_3$)$_3$–(O)C=O–*NH*–), 5.77 (s, 1H, –*CH*=C(CH$_3$)–), 3.11-3.14 ((CH$_3$)$_3$–(O)C=O–NH–*CH$_2$*–), 2.86-2.97 (m, 6H, –NH–(CO)–NH–*CH$_2$*– +–*CH$_2$*–NH–(CO)–NH–), 2.10 (s, 3H, –CH$_3$), 1.21-1.46 (m, 25H, –NH–CH$_2$–*CH$_2$*– + –CH$_2$–CH$_2$–*(CH$_2$)$_2$*–CH$_2$–CH$_2$– + *(CH$_3$)$_3$*–(O)C=O–NH–).

FTIR (ATR): ν (cm$^{-1}$) = 1701 (UPy), 1665 (UPy), 1576 (UPy), 1520 (UPy), 1460, 1355, 1311, 1254.

***Characterization data for compound 3:***

$^1$HNMR (DMSO-d$_6$): δ (ppm) = 9.77 (br.s, 1H, Enol –CH=C(*OH*)–), 7.55-7.72 (br. m, 4H, –CH$_2$–NH(CO)–*NH*– + –CH$_2$–*NH*(CO)–NH– + –C(O)–*NH*(CH$_2$)$_6$*NH*–), 5.79 (s, 1H, –*CH*=C(CH$_3$)–),

3.10-3.14, –CONH–$(CH_2)_5$–*$CH_2$*–NHCO–), 2.94-2.97 (m, 4H,–CONH–*$CH_2$*–$(CH_2)_4$–*$CH_2$*–NHCO–), 2.73-2.80 (m, 2H, $N^+H_3$–*$CH_2$*–$(CH_2)_5$–NHCO–), 2.11 (s, 3H, –$CH_3$), 1.22-1.54 (m, 16H, –$CH_2$–*$(CH_2)_4$*–$CH_2$–).

FTIR (ATR): ν ($cm^{-1}$) = 1710 (UPy), 1670 (UPy), 1580 (UPy), 1520 (UPy), 1462, 1355, 1312, 1258, 1201 (TFA Salt), 1135 (TFA Salt).

***Characterization data for compound 4:***

$^1$HNMR (DMSO-$d_6$): δ (ppm) = 11.64 (m, 1H, –$(CH_3)$C–*NH*–), 9.76 (s, 1H, –$CH_2$–NH(CO)–*NH*–), 7.82 (m, 2H, –$CH_2$–*NH*(CO)–NH–), 7.71-7.95 (m, 2H, –C(O)–*NH*$(CH_2)_6$*NH*–C(O)–), 5.76 (s, 1H, –*CH*=C$(CH_3)$–), 3.33 (m, 3H, *$N^+H_3$*–$(CH_2)_5$–*$CH_2$*–NHCO–), 3.10-3.14, –CONH–$(CH_2)_5$–*$CH_2$*–NHCO–), 2.94-2.97 (m, 4H,–CONH–*$CH_2$*–$(CH_2)_4$–*$CH_2$*–NHCO–), 2.73-2.77 (m, 2H, $N^+H_3$–*$CH_2$*–$(CH_2)_5$–NHCO–), 2.10 (s, 3H, –$CH_3$), 1.22-1.56 (m, 16H, –$CH_2$–*$(CH_2)_4$*–$CH_2$–).

FTIR (ATR): ν ($cm^{-1}$) = 1699 (UPy), 1669 (UPy), 1575 (UPy), 1520 (UPy), 1466, 1357, 1312, 1256.

#### Supplementary Methods

***Rat Joint OARSI Scoring.*** Extent of degeneration was determined as described by Gerwin et al (*1*). with a few modifications. Two forms of assessments were performed: semi-quantitative joint scoring and quantitative measurements of degeneration. For scoring, there were four areas of evaluation, including total tibial cartilage degeneration (0-5 for 3 zones, total 0-15), femoral cartilage degeneration (0-5), bone score (0-5), and osteophyte score (0-4), with total added score ranging from 0-29. Tibial and femoral cartilage degeneration was assessed based upon the percentage of total cartilage with matrix fibrillation or matrix loss or chondrocyte death, but not proteoglycan loss, using the scoring criteria described in Gerwin et al (*1*). For evaluating tibial cartilage degeneration, the total width of the cartilage was separated into three approximate regions and scored independently. For the femur, the total cartilage width was evaluated as a whole. The bone score was assessed exactly as described by evaluating the subchondral bone below the most severe cartilage lesion(*1*). A total of four sections, two from two separate regions in the medial compartment of the joint, were evaluated by three blinded independent scorers. For scores that differed by more than 2.5 from the average for tibial cartilage degeneration, 1.5 from the average for femoral cartilage degeneration, and 2 from the average for bone score, joints were rescored so that consensus was reached amongst all scorers. The osteophyte score was modified for sagittal joint sections based upon a histogram of osteophyte sizes for all samples in the mi-ACLT groups (saline, HA, and HA-UPy). The modified osteophyte score is shown in Table S1. Osteophyte size was measured for each of the four sections using ImageJ. An average of the total joint score for the four sections was reported as one score per subject.

**Table S1.** Scoring criteria for osteophyte evaluation. Modified from Gerwin et al.(*1*)

| Grade | Description |
|---|---|
| 0 | No osteophytes |
| 1 | < 299 μm |
| 2 | 300-599 μm |
| 3 | 600-899 μm |
| 4 | > 900 μm |

Measurements of cartilage degeneration were performed using ImageJ as described (*1*). Parameters of cartilage degeneration that were assessed included total and significant cartilage degeneration, surface matrix loss, and zonal depth ratio of lesions. As a modification, these parameters were evaluated for both the tibia and the femur. Total and significant cartilage degeneration and surface matrix loss were expressed as a percentage of the total width of the cartilage. Total cartilage degeneration was evaluated as the width of cartilage with type of degeneration, including matrix loss or fibrillation, or proteoglycan loss with or without chondrocyte death. Significant cartilage degeneration was evaluated as the width of cartilage where 50% or more of the cartilage thickness from surface to tidemark shows matrix loss or chondrocyte loss or death. Surface matrix loss included all collagen matrix loss at 0% depth of the cartilage, particularly observed as fibrillation. It does not evaluate proteoglycan loss or chondrocyte death. The zonal depth ratio of lesions was evaluated as the ratio of the total cartilage thickness that shows any type of cartilage degeneration, measured at the midpoint of each of the three zones of the cartilage. The reported value is an average of the three zones. The measurements were carried out by one individual blinded to the groups. As with the semi-quantitative score, a total of four sections were evaluated, and the reported values represented an average of the four sections for each subject.
